# Replicable Connectivity and Spectral Alterations of Functional Networks in Major Depression

**DOI:** 10.64898/2026.11.06.764930

**Authors:** Bradley Baker, Malvika Sridhar, Vince Calhoun

**Affiliations:** Tri-Institutional Center for Translational Research in Neuroimaging and Data Science (TReNDS), Georgia State University, Georgia Institute of Technology, and Emory University

## Abstract

Major depressive disorder (MDD) is usually characterized in functional MRI through connectivity alone, often in samples too small to resolve its modest effects. Here we applied multi-scale, template-guided independent component analysis to resting-state data from the multi-site MDD-DIRECT cohort and jointly examined the connectivity, spectral content and spatial composition of 105 intrinsic brain networks. Differences between patients and controls were widespread, small and structured: cerebello-subcortical coupling was reduced, subcortical–somatomotor coupling was increased, and spectral power shifted from lower to higher frequencies, most prominently in the thalamus, which also showed the most altered connectivity. Spatial composition differed in every network examined. In a smaller independent cohort, the direction of effect agreed for 95% of significant connections, 78% of significant spectral effects and 77% of the largest spatial-composition effects. Depression therefore alters several distinct properties of intrinsic networks, consistently across cohorts and most prominently in the thalamus.

## Introduction

With over 180 million cases as of 2019 [1, 2], major depressive disorder (MDD) is the most prevalent mood disorder across the globe. More recently, MDD continues to increase in prevalence, seemingly spurred on in part by global events such as the COVID-19 pandemic in 2020 [3]. In 2022, the World Health Organization named Depression as the second highest burden and disability causing disease among all diseases in 2020, and projects it to become the largest burden globally by 2030 [4]. On an individual level, MDD is frequently debilitating, with patients experiencing depressed mood, loss of interest or pleasure, significant weight loss or gain, sleep disturbance, psychomotor agitation, tiredness and fatigue, impaired cognition, and recurrent thoughts of death, suicidal ideation or attempts[5]. This combination of systemic and individual suffering has led to increased attention from policy-makers and researchers worldwide [6], spurring efforts to further understand the underlying neurological mechanisms which are affected in MDD, associated with symptom severity, and treatment outcomes.

Neuroimaging provides a mechanism for the study of the neurological bases of MDD, with the goal not only to further understanding the underlying mechanisms, but to also eventually inform clinical intervention or diagnosis via quantifiable biomarkers. Structural markers from structural and diffusion imaging such as decreased gray and white matter volume in the hippocampus, cerebellum, and other regions [7, 8, 9, 10, 11, 12] have been widely identified in the literature. Numerous EEG studies have pinpointed abnormalities in connectivity, power differences across frequency bands in patients, and other oscillatory biomarkers[13, 14, 15, 16]. Functional magnetic resonance imaging (fMRI), which measures the blood-oxygen-level-dependent (BOLD) signal associated with increased neural activity, provides an important complementary component in the study of MDD, allowing unique insights into functional changes within the brain. Importantly, fMRI provides insight into whole brain activity, and can help identify functional alterations not accessible to EEG, for example.

Many fMRI-based MDD biomarkers have been proposed in the literature. One of the most common is functional connectivity (FC), in which voxel- or ROI-level BOLD activation is correlated over the period of the scanning session, providing insight into co-activation between regions. FC has been widely identified as “abnormal” in MDD, with studies often characterizing MDD as a disorder of functional dys-connectivity [17]. Specific regional FC findings tend to differ between studies, though a number have identified abnormal FC particularly in the saliency and default mode networks. Functional network connectivity (FNC) is a related, but methodologically distinct measure of brain connectivity, which uses the time-series from functional networks derived from data using methods like independent component analysis (ICA) [18], rather than fixed ROIs or voxelwise connectivity. Studies of MDD with FNC have been comparatively limited, but findings have similarly revealed widespread altered connectivity in MDD [19]. Our work contributes to this growing body of literature, complementing previous studies of FNC in MDD, extending beyond connectivity biomarkers to present a more holistic picture of functional alterations in MDD which studies frequency-domain and alterations in functional network size in addition to connectivity.

Although the limited temporal resolution in fMRI limits the scope of frequency-based analyses, the marriage of time-varying BOLD signal with a higher spatial resolution than EEG allows fMRI to provide unique, region-specific insights into the spectral characteristics of MDD. The majority of historical fMRI studies of MDD have reported alterations in aggregate measures of frequency content, such as fractional amplitude of low-frequency fluctuations (fALFF). Alterations in fALFF have been consistently identified within the cerebellum [20, 21, 22, 23, 24], as well as within the parahippocampal gyrus[21, 22, 25, 26], right middle frontal gyrus [20, 22], precuneus and cuneus [22, 25, 27, 24] and putamen [20, 25]. Other novel aggregated measures such as the power spectral slope, which captures both high and low amplitude fluctuations, suggests that MDD patients show a sharper change in power between high and low frequencies[28]. Comparatively fewer studies have provided non-aggregate characterizations of fMRI spectra, with focus given to differences in determining focal distributions of high-frequency activity[29, 30] such as the saliency network, and development of frequency-resolved connectivity profiles[31] across the brain. Although the literature is varied in terms of particular findings, differences in BOLD frequency content have been reported consistently enough to suggest that the frequency composition of the signal carries information about MDD which band-limited summary measures such as fALFF do not fully capture. What has been missing is a characterization of how that content is distributed across the spectrum, in a cohort large enough to resolve effects of the size the disorder appears to produce.

Some recent studies have focused on the spatial and topographical characteristics of particular functional networks. One recent work for example, characterized MDD by a distinct expansion of the saliency functional network [32]. Altogether, however, since many fMRI studies are tied to fixed, atlas-based ROIs, work along this axis has been relatively limited. Our analysis in this work thus represents a unique contribution via its use of the NeuroMark ICA method [33, 34], which can support variation in functional network size based on characteristics of the data, while still anchoring findings to well-established resting-state networks found across large populations.

In this work, we present a comprehensive study of the large-scale (N=3505) MDD-DIRECT fMRI data set[35], contributing to the growing body of work uncovering neuroimaging biomarkers for major depression. In particular, our analysis methodology is based on the NeuroMark ICA framework [33, 34], which bridges the gap between data-driven specificity and template-driven replicability. Our contributions expand the growing body of work characterizing functional network connectivity differences in MDD, and offer a rich characterization of spectral features as of yet widely unexplored in large studies. Additionally, we assess the cross-cohort consistency of these features in a smaller, independently collected cohort from Zhi et al. [19], comparing the direction of effects network by network, a direct comparison that template-driven methods make possible without post hoc matching of components.

## Results

The 105 intrinsic-connectivity networks (ICNs) identified using the NeuroMark ICA 2.2 template[34] are provided in Fig. 1. Each ICN is categorized into one of 7 domains: Cerebellum (CB), Visual (VI), Paralimbic (PL), Subcortical (SC), Somatomotor (SM), Higher Cognitive (HC) and Triple Network (TN). The visual domain is further divided into the OT and OC subdomains; the subcortical domain is divided into the EH, ET, and BG subdomains; the higher cognitive (HC) domain is divided into the IT, TP, and FR subdomains, and the triple network (TN) domain is divided into the CE, Default Mode (DM) and Saliency (SA) subdomains.

**Fig. 1.**
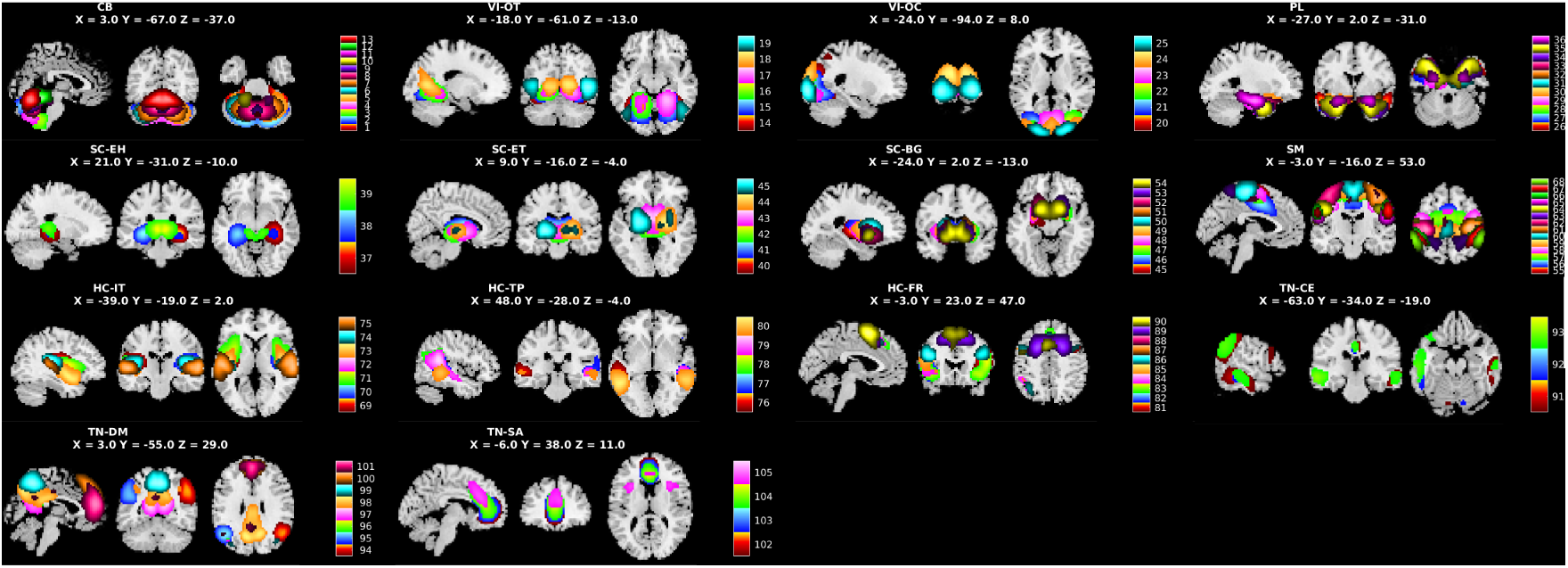
A domain-level view of the 105 networks contained in the NeuroMark2.2 template [34].

We begin our study by examining bulk cross-sectional differences between controls and individuals with MDD, and we then proceed to extend this analysis to the study of symptom severity by looking at the total symptom scores (HAMDTotal, HAMATotal). All diagnostic tests are encoded as MDD-Control tests so that negative *T* scores are associated with decreased feature measures in MDD and positive *T* scores associated with increases in MDD. Similarly, for symptom score tests, negative *T* values indicate that higher symptom scores are associated with feature decreases and positive *T* values indicate that higher symptom scores are associated with increased feature measures.

### Diagnosis

We first study the differences between MDD patients and controls to broadly characterize alterations in static functional network connectivity (sFNC), timecourse spectra, and spatial map composition.

### Static FNC

We observe widespread alterations in sFNC associated with diagnosis in the MDD-DIRECT data. Fig. 2 presents the sFNC findings in MDD-DIRECT associated with MDD diagnosis. Panel 2a shows a chord diagram with the top 1% of total connections (55/5460) sorted by Cohen’s *d* effect size. 651 of 5460 connections (11.9%) were significant, of which 379 (58%) reflected reduced connectivity in MDD and 272 (42%) increased connectivity. MDD in this cohort is therefore characterized by a mixture of hypo- and hyper-connectivity rather than a uniform shift in either direction. Effect sizes were small throughout: the largest was *|d|* = 0.403, 276 of the 651 significant connections reached at least a minor effect (*|d| ≥* 0.2), and only 8 reached *|d| ≥* 0.3. Alterations were preferentially inter-domain rather than intra-domain, affecting 12.1% of between-domain pairs (612/5039) but only 9.3% of within-domain pairs (39/421), indicating that, in this sample, MDD manifests principally as disrupted coupling between functional systems rather than the internal coherence of individual systems. Tables 1 and 2 summarize the significant connections by domain. The group mean sFNC matrices and their difference are shown in Supplementary Note 1 (Supplementary Fig. 1).

**Fig. 2.**
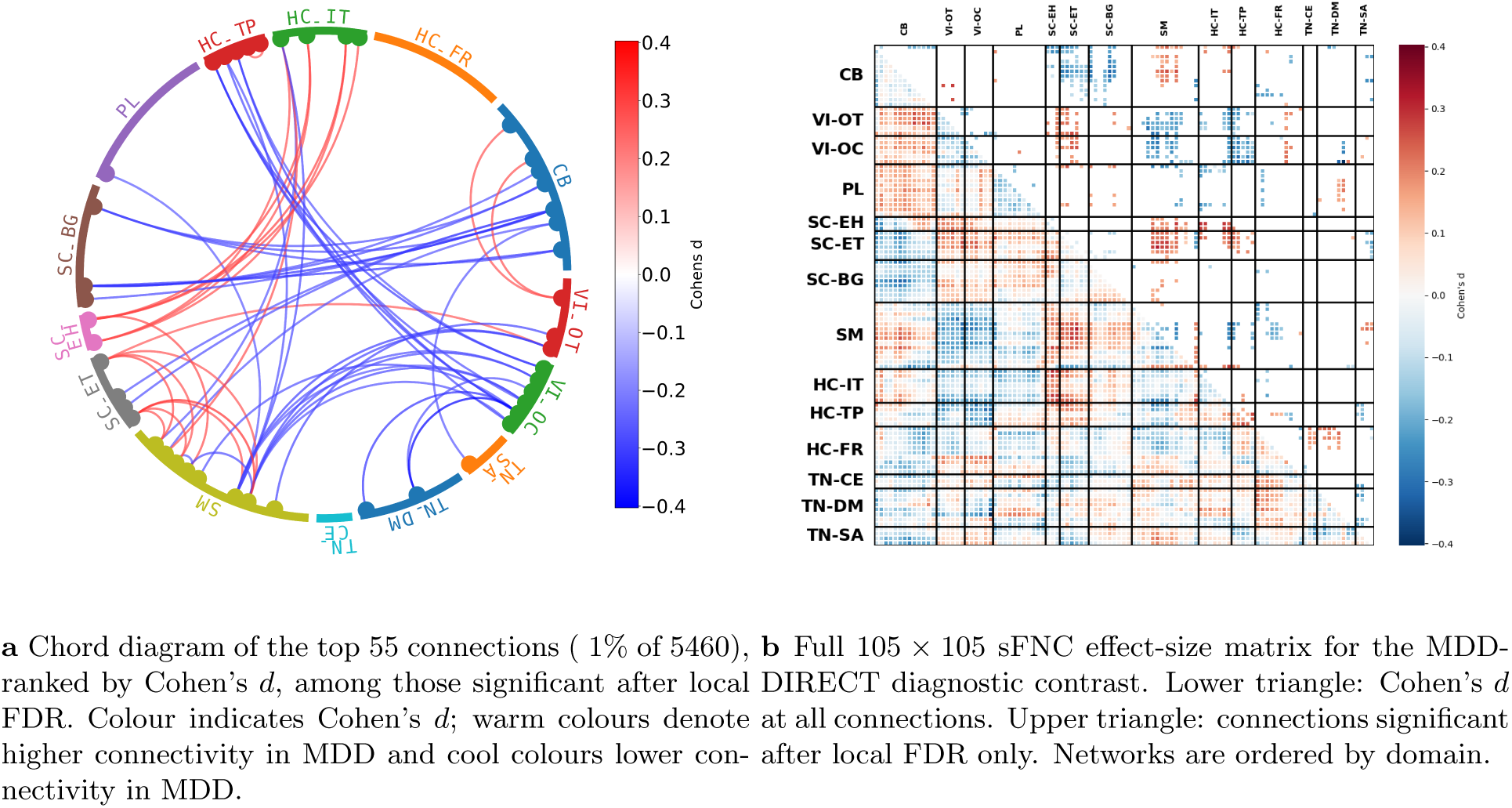
Static functional network connectivity differences between MDD patients and controls in MDD-DIRECT. *n* = 1,123 patients with MDD and 831 controls. Each connection was compared between groups by ANCOVA adjusting for age, sex and site, with local FDR correction (*q ≤* 0.05) across all 5,460 connections; effect size is Cohen’s *d*.

**Table 1.** Connections differing significantly between MDD patients and controls in MDD-DIRECT, by domain (local FDR *q ≤* 0.05; 651 connections in total). A connection is counted under each of its two domains. *N* ^+^ and *N ^−^*: connections with higher or lower connectivity in MDD. ANCOVA adjusting for age, sex and site (*n* = 1,123 patients with MDD and 831 controls); effect size is Cohen’s *d*; Min *p* is uncorrected.

| Domain | Total connections | Mean \|*d*\| | Max \|*d*\| | Min *p* | *N*<sup>+</sup> | *N*<sup>−</sup> |
|---|---|---|---|---|---|---|
| CB | 152 | 0.193 | 0.321 | 1.36e-08 | 45 | 107 |
| VI-OT | 99 | 0.198 | 0.300 | 2.52e-08 | 33 | 66 |
| VI-OC | 103 | 0.206 | 0.403 | 1.54e-11 | 27 | 76 |
| PL | 63 | 0.179 | 0.263 | 1.32e-10 | 27 | 36 |
| SC-EH | 48 | 0.196 | 0.298 | 1.55e-08 | 38 | 10 |
| SC-ET | 152 | 0.185 | 0.321 | 1.32e-10 | 89 | 63 |
| SC-BG | 68 | 0.180 | 0.321 | 1.36e-08 | 23 | 45 |
| SM | 230 | 0.200 | 0.321 | 7.36e-10 | 112 | 118 |
| HC-IT | 79 | 0.203 | 0.298 | 1.32e-07 | 31 | 48 |
| HC-TP | 84 | 0.195 | 0.310 | 1.54e-11 | 27 | 57 |
| HC-FR | 89 | 0.179 | 0.255 | 2.67e-08 | 38 | 51 |
| TN-CE | 13 | 0.171 | 0.228 | 1.16e-06 | 8 | 5 |
| TN-DM | 44 | 0.188 | 0.403 | 4.25e-11 | 26 | 18 |
| TN-SA | 39 | 0.182 | 0.265 | 3.36e-07 | 8 | 31 |

**Table 2.** Connections differing significantly between MDD patients and controls in MDD-DIRECT (local FDR. *q ≤* 0.05**), by pair of domains.** Diagonal entries are within-domain connections. ANCOVA adjusting for age, sex and site (*n* = 1,123 patients with MDD and 831 controls).

| Domain | CB | VI-OT | VI-OC | PL | SC-EH | SC-ET | SC-BG | SM | HC-IT | HC-TP | HC-FR | TN-CE | TN-DM | TN-SA |
|---|---|---|---|---|---|---|---|---|---|---|---|---|---|---|
| CB | 7 | 3 | 0 | 2 | 9 | 40 | 32 | 32 | 7 | 2 | 11 | 0 | 2 | 5 |
| VI-OT | 3 | 2 | 6 | 1 | 8 | 14 | 3 | 30 | 14 | 12 | 6 | 0 | 0 | 0 |
| VI-OC | 0 | 6 | 1 | 2 | 4 | 13 | 0 | 28 | 7 | 27 | 7 | 0 | 6 | 2 |
| PL | 2 | 1 | 2 | 3 | 1 | 5 | 7 | 9 | 15 | 0 | 6 | 1 | 11 | 0 |
| SC-EH | 9 | 8 | 4 | 1 | 1 | 0 | 0 | 16 | 8 | 1 | 0 | 0 | 0 | 0 |
| SC-ET | 40 | 14 | 13 | 5 | 0 | 3 | 9 | 37 | 10 | 13 | 0 | 0 | 1 | 7 |
| SC-BG | 32 | 3 | 0 | 7 | 0 | 9 | 0 | 14 | 1 | 0 | 2 | 0 | 0 | 0 |
| SM | 32 | 30 | 28 | 9 | 16 | 37 | 14 | 15 | 10 | 11 | 20 | 0 | 2 | 6 |
| HC-IT | 7 | 14 | 7 | 15 | 8 | 10 | 1 | 10 | 1 | 2 | 4 | 0 | 0 | 0 |
| HC-TP | 2 | 12 | 27 | 0 | 1 | 13 | 0 | 11 | 2 | 4 | 6 | 0 | 0 | 6 |
| HC-FR | 11 | 6 | 7 | 6 | 0 | 0 | 2 | 20 | 4 | 6 | 2 | 10 | 13 | 2 |
| TN-CE | 0 | 0 | 0 | 1 | 0 | 0 | 0 | 0 | 0 | 0 | 10 | 0 | 0 | 2 |
| TN-DM | 2 | 0 | 6 | 11 | 0 | 1 | 0 | 2 | 0 | 0 | 13 | 0 | 0 | 9 |
| TN-SA | 5 | 0 | 2 | 0 | 0 | 7 | 0 | 6 | 0 | 6 | 2 | 2 | 9 | 0 |

We also observe that significantly altered sFNC is not distributed uniformly across domains. The somatomotor (SM, 230 connections), cerebellar (CB, 152) and extended thalamus (SC-ET, 152) domains are the most heavily involved in connectivity differences, and SC-ET shows the highest proportion of altered connections of any domain (152/609, 25.0%). Conversely, the three triple-network domains are the least affected, both absolutely and proportionally: central executive (TN-CE, 13/309), default mode (TN-DM, 44/804) and salience (TN-SA, 39/410). While this does not contradict the prevalence of these networks in the connectivity literature, it does suggest that the whole-brain effects may be dominated by other alterations.

Four blocks of connections account for a large share of the total and are notable for being almost entirely consistent in direction between individual connectivity pairs. First, connectivity between the cerebellum and the subcortical domains is uniformly reduced in MDD: CB–SC-BG (32 connections), CB–SC-ET (40) and CB–SC-EH (9) are uniformly lower in MDD. The strongest of these are between left cerebellum and left caudate (*d* = *−*0.321, *p* = 1.4 *×* 10*^−^*^8^), bilateral cerebellum and left caudate (*d* = *−*0.312), and cerebellum and dorsal striatum/thalamus (*d* = *−*0.285). Second, and in the opposite direction, connectivity between the subcortical domains and somatomotor cortex is almost uniformly increased in MDD: SC-ET–SM (37 connections), SC-EH–SM (16) and SC-BG–SM (14), with the largest effects between anterior diencephalon and precentral gyrus (*d* = +0.321) and between diencephalon/midbrain and paracentral lobule (*d* = +0.285). Third, connectivity between visual and temporal or default-mode networks is uniformly reduced in MDD, most strikingly between the occipital-calcarine and temporal-parietal subdomains (VI-OC–HC-TP, 27 of the 30 possible connections in that block, all negative). This block contains both the largest effect in the analysis, between occipital pole and dorsal precuneus (*d* = *−*0.403, *p* = 4.3 *×* 10*^−^*^11^), and the most significant, between occipitotemporal junction and superior temporal gyrus (*p* = 1.5 *×* 10*^−^*^11^, *d* = *−*0.310). Finally, the extended hippocampal domain (SC-EH) is the most directionally lopsided in the study, with 38 increased against 10 decreased connections, driven by increased coupling between hippocampal, parahippocampal and amygdalar networks and inferior parietal and postcentral cortex (peak *d* = +0.298).

At the level of individual networks, the most frequently implicated are precentral gyrus (37 significant connections of 104 possible), thalamus/hippocampus (35), left superior postcentral gyrus (35), superior sensorimotor cortex (33), paracentral lobule (33) and posterior supplemental motor area (32).

We next assessed whether these connectivity differences are consistent in the smaller, independently collected cohort of Zhi et al. [19]. Because that cohort has limited power for effects of the size observed here (Methods), we assess consistency by the direction of effect rather than by significance. Of the 651 connections significant in MDD-DIRECT, 620 (95.2%) show the same direction of effect in Zhi et al., against 50% expected if the two cohorts were unrelated (one-sided binomial *p <* 10*^−^*^100^). Agreement is 96.0% (265 of 276) for connections with *|d| ≥* 0.2, 53 of the 55 connections shown in Fig. 2a, and 8 of 8 with *|d| ≥* 0.3. It is near-complete in each of the blocks described above: 75 of 81 cerebello-subcortical connections, 65 of 67 subcortical–somatomotor connections and 27 of 27 VI-OC–HC-TP connections agree in direction. Across all 5460 connections, irrespective of significance, the direction agrees in 74.8% and the unthresholded *T* maps correlate at *r* = 0.67. The Zhi et al. results and the overlap of significant connections between cohorts are given in Supplementary Note 2 and Supplementary Fig. 2.

### Spectra

Spectral power was compared between groups within each of 129 frequency bins for each of the 105 networks. 432 of 13,545 bins (3.2%) differed significantly between MDD patients and controls, 234 reflecting increased and 198 decreased power in MDD (Tables 3 and 4).

**Table 3.** Spectral bins differing significantly between MDD patients and controls in MDD-DIRECT, by domain (local FDR *q ≤* 0.05; 432 bins in total). ANCOVA in each frequency bin adjusting for age, sex and site (*n* = 1,123 patients with MDD and 831 controls); effect size is Cohen’s *d*; Min *p* is uncorrected.

| Domain | Significant bins | Mean \|*d*\| | Max \|*d*\| | Min *p* |
|---|---|---|---|---|
| CB | 48 | 0.155 | 0.188 | 1.20e-05 |
| VI-OT | 12 | 0.159 | 0.194 | 2.54e-03 |
| VI-OC | 7 | 0.161 | 0.193 | 1.25e-03 |
| PL | 18 | 0.143 | 0.216 | 8.53e-05 |
| SC-EH | 8 | 0.175 | 0.237 | 6.35e-05 |
| SC-ET | 99 | 0.137 | 0.210 | 2.87e-06 |
| SC-BG | 78 | 0.173 | 0.267 | 2.03e-06 |
| SM | 61 | 0.180 | 0.231 | 3.13e-05 |
| HC-IT | 34 | 0.146 | 0.185 | 3.72e-05 |
| HC-TP | 3 | 0.133 | 0.147 | 3.28e-03 |
| HC-FR | 3 | 0.093 | 0.143 | 3.14e-03 |
| TN-CE | 1 | 0.121 | 0.121 | 4.51e-03 |
| TN-DM | 45 | 0.178 | 0.235 | 6.85e-05 |
| TN-SA | 15 | 0.197 | 0.232 | 4.09e-04 |

**Table 4.** Spectral bins differing significantly between MDD patients and controls in MDD-DIRECT, by frequency range (local FDRq ≤ 0.05; 432 bins in total). ANCOVA in each frequency bin adjusting for age, sex and site (*n* = 1,123 patients with MDD and 831 controls); effect size is Cohen’s *d*; Min *p* is uncorrected.

| Frequency range | Significant bins | Mean \|*d*\| | Max \|*d*\| | Min *p* |
|---|---|---|---|---|
| 0.0000-0.0500 Hz | 118 | 0.171 | 0.267 | 2.87e-06 |
| 0.0500-0.1000 Hz | 85 | 0.148 | 0.189 | 1.20e-05 |
| 0.1000-0.1500 Hz | 74 | 0.156 | 0.235 | 1.31e-05 |
| 0.1500-0.2000 Hz | 96 | 0.152 | 0.232 | 2.03e-06 |
| 0.2000-0.2500 Hz | 59 | 0.173 | 0.214 | 3.13e-05 |

Power is reduced in MDD at low frequencies and increased at high frequencies, with the crossover falling within the conventional resting-state band. Averaged across all networks, the mean *T* statistic is *−*0.258 below 0.010 Hz and *−*1.161 in the 0.010–0.027 Hz band, where only 17.9% of bins show positive *T* ; it rises through zero across 0.027–0.098 Hz and reaches +0.920 above 0.150 Hz, where 84.6% of bins are positive. Among the significant bins, those showing reduced power in MDD are centred at 0.029 Hz and those showing increased power at 0.168 Hz. MDD is therefore associated within the cohort with a shift of spectral power from lower to higher frequencies, that is, a flattening of the BOLD power spectrum. The same shift is visible in the spectrum averaged across all 105 networks (Fig. 3): averaged power in MDD is lower than in controls around the low-frequency peak (approximately 0.01–0.03 Hz) and slightly higher at the highest frequencies, although for this averaged spectrum the adjusted *T* statistic exceeds the uncorrected *p* = 0.05 threshold only in narrow bands. Fig. 4 shows the differences for each network and frequency bin.

**Fig. 3.**
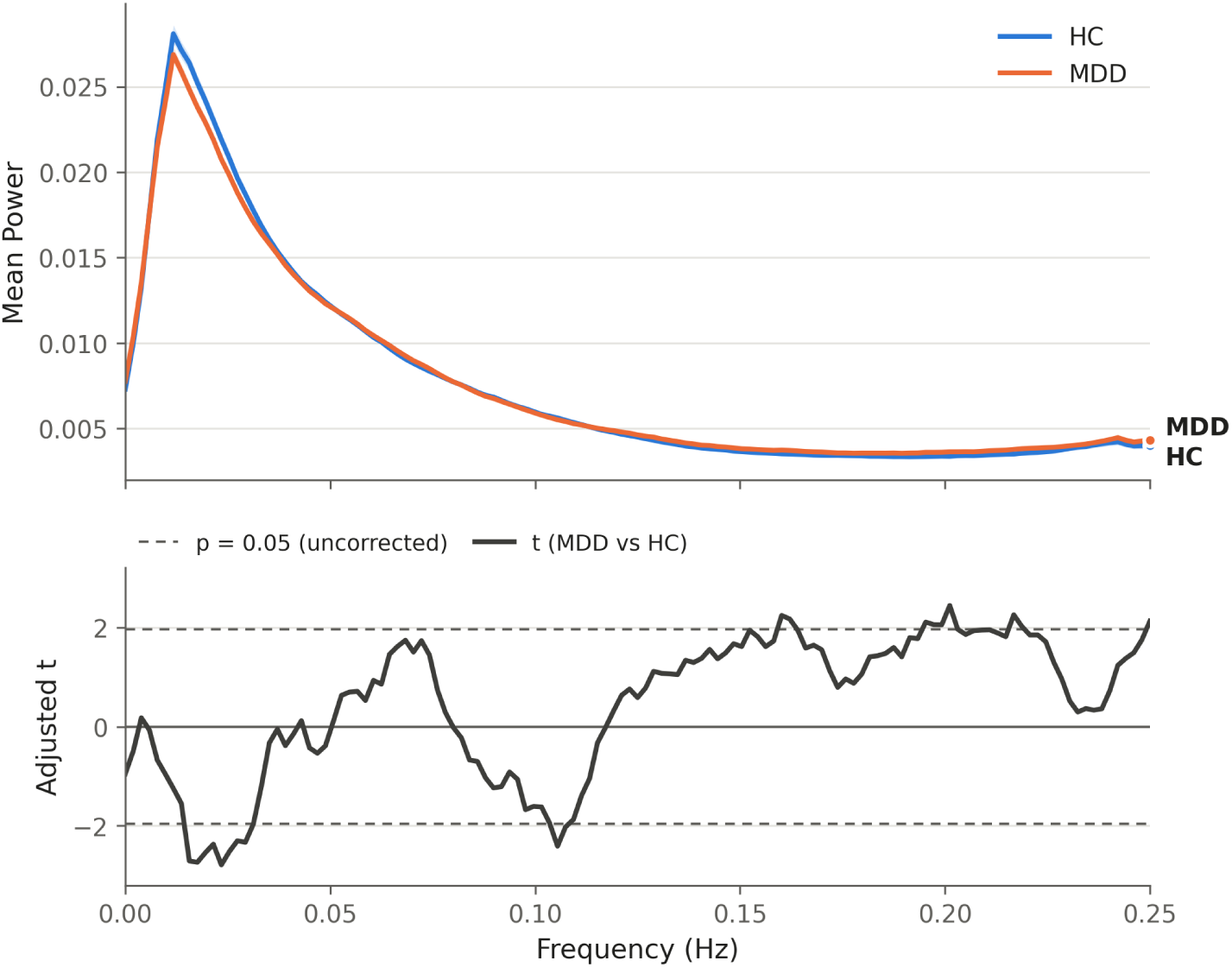
Spectral power averaged across all 105 networks in MDD-DIRECT. Top, mean power spectrum of controls (HC, blue line; *n* = 831) and MDD patients (orange line; *n* = 1,123), averaged over networks and participants;. Bottom, covariate-adjusted *T* statistic for MDD versus controls at each frequency (solid line); dashed lines mark the uncorrected two-sided *p* = 0.05 threshold. Power is lower in MDD around the low-frequency peak and slightly higher at the highest frequencies.

**Fig. 4.**
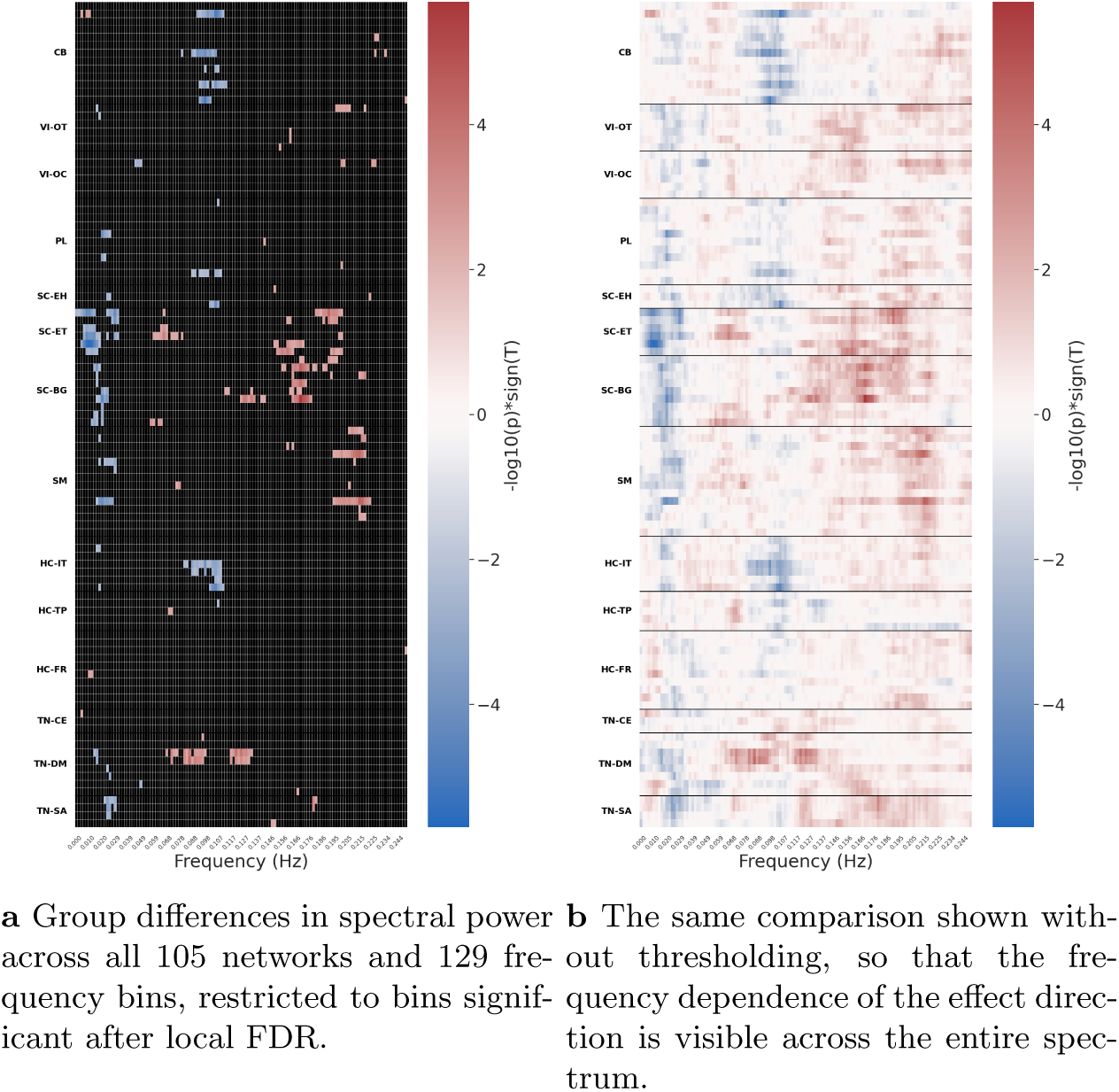
Spectral power differences between MDD patients and controls in MDD-DIRECT. Power is reduced in MDD at low frequencies and increased at high frequencies. *n* = 1,123 patients with MDD and 831 controls. Power in each of 129 frequency bins of each of 105 networks was compared between groups by ANCOVA adjusting for age, sex and site, with local FDR correction (*q ≤* 0.05) across all 13,545 network–frequency tests; effect size is Cohen’s *d*.

Networks in the extended thalamus domain (SC-ET) carry the largest share of significant bins, and within these networks, the frequency dependence is explicit: right thalamus, anterior diencephalon and diencephalon/midbrain show reductions confined to 0.004–0.031 Hz, while left thalamus and thalamus/hippocampus show increases at 0.152–0.201 Hz. The strongest individual reductions are in right thalamus at 0.0098 Hz (*T* = *−*4.69, *p* = 2.9 *×* 10*^−^*^6^, *d* = *−*0.194) and thalamus/hippocampus at 0.0078 Hz (*d* = *−*0.176); the strongest increase is in right caudate at 0.170 Hz (*T* = +4.77, *d* = +0.226). Beyond the thalamus, the cerebellum (dorsomedial and posterior) and the insula (right, anterior, and medial ventral anterior) show reduced power near 0.10 Hz.

In the Zhi et al. cohort, 337 of the 432 bins significant in MDD-DIRECT (78.0%) show the same direction of effect (one-sided binomial *p* = 4 *×* 10*^−^*^33^), as do 35 of the 48 with *|d| ≥* 0.2 (72.9%). Agreement is high below 0.150 Hz (83.8% below 0.027 Hz, 83.7% across 0.027–0.098 Hz and 92.5% across 0.098–0.150 Hz) and in the extended thalamus (86.9%), cerebellum (91.7%) and default mode (93.3%), but weaker above 0.150 Hz (62.7%) and at chance in the basal ganglia (52.6%). Across all 13,545 bins the direction agrees in 59.9%, and the unthresholded *T* maps correlate at *r* = 0.31. The Zhi et al. spectra and the overlap of significant bins are given in Supplementary Note 2 and Supplementary Fig. 3.

### Spatial Maps

Finally, we compared the intensity of each participant’s estimated spatial maps voxelwise between groups. Local FDR alone leaves a very large number of voxels (2,219,040, or 29.6%), but the median effect size among them is *|d|* = 0.076, well below any conventional threshold for interpretability. Restricted to each network’s template extent (*|Z| >* 2), 79,850 voxel–network tests survive local FDR, 1,886 of them with *|d| ≥* 0.2 and only one with *|d| ≥* 0.3; Table 5 gives the number of distinct significant voxels within each domain. Fig. 5 shows significant voxels within the template extents thresholded by effect size *|d| >* 0.2 to emphasize the largest effects. The results indicate that spatial map differences in MDD are present in every domain, and that domains differ in the proportion of their extent affected rather than in whether they are affected at all.

**Fig. 5.**
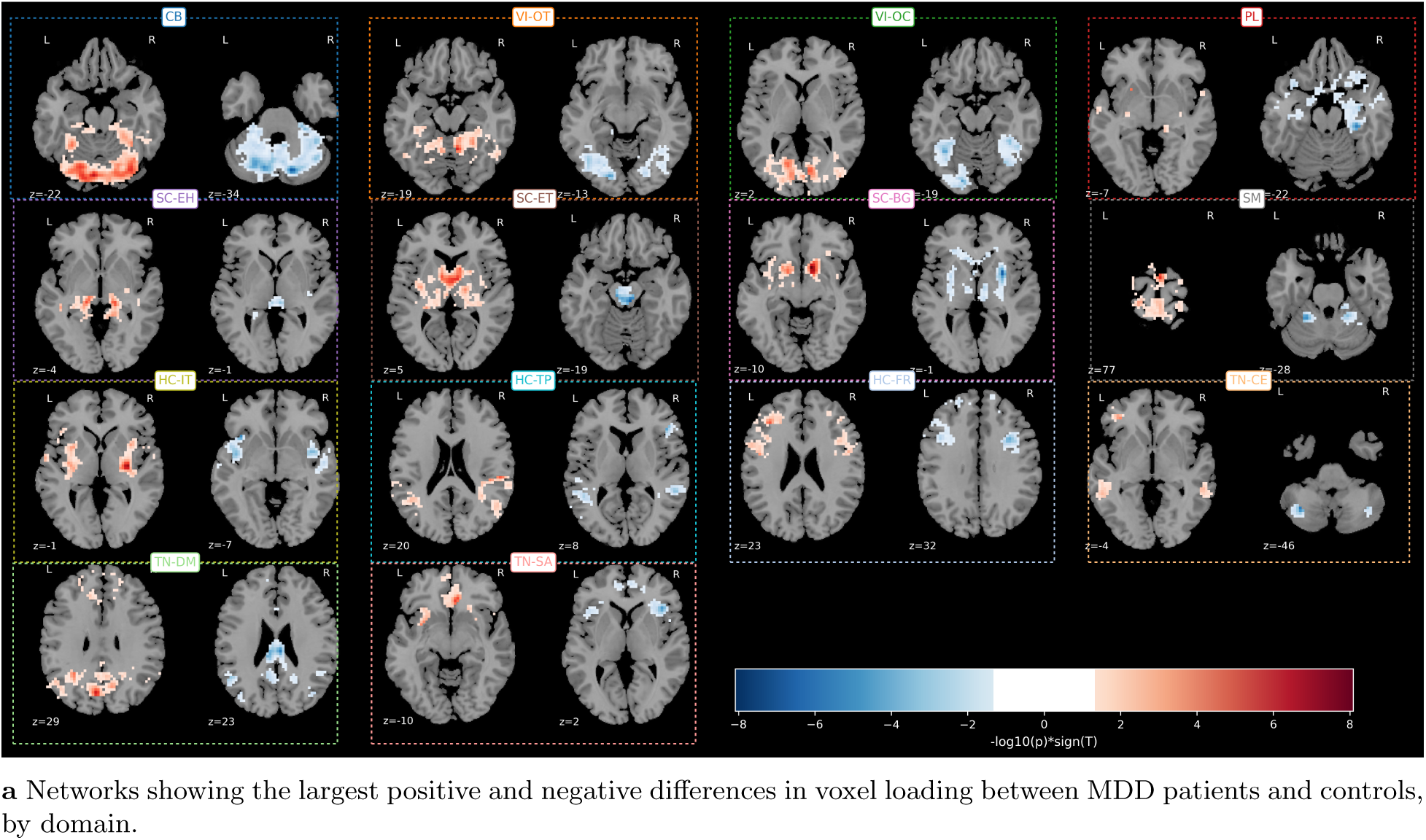
Spatial map differences between MDD patients and controls in MDD-DIRECT. Positive values denote higher voxel loading on that network in MDD. *n* = 1,123 patients with MDD and 831 controls. Voxel loadings were compared between groups by ANCOVA adjusting for age, sex and site, with local FDR correction (*q ≤* 0.05) across voxels and networks. Voxels are restricted to each network’s NeuroMark 2.2 template extent (*|Z| >* 2) and shown where significant after local FDR (*q ≤* 0.05) with *|d| ≥* 0.2; colour shows Cohen’s *d*.

**Table 5.** Number of distinct voxels significant after local FDR (*q ≤* 0.05) for the diagnosis contrast, by domain, with each network restricted to its NeuroMark 2.2 template extent. (*|Z| >* 2). Voxelwise ANCOVA adjusting for age, sex and site (*n* = 1,123 patients with MDD and 831 controls).

| Domain | Significant voxels |
|---|---|
| CB | 6438 |
| VI-OT | 3453 |
| VI-OC | 4325 |
| PL | 3344 |
| SC-EH | 1120 |
| SC-ET | 2045 |
| SC-BG | 2763 |
| SM | 8375 |
| HC-IT | 3703 |
| HC-TP | 2620 |
| HC-FR | 5591 |
| TN-CE | 2454 |
| TN-DM | 6213 |
| TN-SA | 2259 |

Looking at only the largest (but still relatively weak) effects (*|d| >* 0.2), the cerebellum contains the largest share of surviving voxels (1,730 across 13 ICNs) and is dominated by increased spatial map weighting in MDD (1,138 increased against 592 decreased), followed by the somatomotor domain (733 voxels across 14 networks), which is narrowly inclined toward a decrease in MDD (246 against 487). The occipital-calcarine and occipitotemporal visual domains contribute 487 and 469 voxels, the default mode 443, and the extended thalamus 442; the basal ganglia are strongly increase-dominant (293 voxels, 237 increased against 56 decreased) and the central executive domain strongly decrease-dominant (135 voxels, 29 against 106). At the stricter criterion of *|d| ≥* 0.25, 464 voxels in 56 networks remain, of which the cerebellum accounts for 206 across 10 networks. The networks contributing most are dorsomedial cerebellum (332 voxels, peak *d* = +0.318), right and left anterior dorsomedial cerebellum (262 and 213), anterior dorsomedial cerebellum (217, peak +0.296), posterior cerebellum (189), striate cortex (159), fusiform gyrus (157), cuneus (152), vermis (149), superior sensorimotor cortex (132, peak *−*0.285), anterior diencephalon (127, peak +0.287) and left superior postcentral gyrus (125). Within the template extents the ordering is the same: the cerebellum supplies 743 of the 1,886 voxels at *|d| ≥* 0.2, followed by the extended thalamus (194) and the basal ganglia (124).

In the Zhi et al. cohort the direction of effect agrees at 77.2% (4,912 of 6,364) of the voxels meeting the primary criterion in MDD-DIRECT (local FDR with *|d| ≥* 0.2), rising to 81.7% (379 of 464) at *|d| ≥* 0.25 and falling to 66.3% at *|d| ≥* 0.1 and 56.0% across all voxels. At the primary criterion, agreement ranges across domains from 70.6% in the basal ganglia to 82.4% in the extended hippocampal domain. The unthresholded effect maps, however, correlate only weakly (*r* = 0.15). The Zhi et al. spatial maps and their within-cohort analysis are given in Supplementary Note 2 and Supplementary Fig. 4.

### Symptom Severity

We next examine whether the same derivatives track symptom severity within the patient group, using total scores on the Hamilton Anxiety (HAMA) and Hamilton Depression (HAMD) rating scales. Severity scores were available for a subset of the cohort, giving effective sample sizes of approximately 663 participants for HAMA and 925 for HAMD, against approximately 1,930 total for the diagnostic contrast. Because the target variables are continuous, we report the partial correlation between score and connectivity after adjustment for covariates as the effect size.

### Static FNC

Associations with severity were considerably sparser and weaker than the diagnostic differences. 246 of 5460 connections (4.5%) were significantly associated with HAMA total score, 139 positively and 107 negatively, and 134 (2.5%) with HAMD total score, 72 positively and 62 negatively (Figs. 6 and 7; Tables 6–9). The largest partial correlations were 0.122 for HAMA and 0.114 for HAMD, well below the effect sizes seen in the diagnostic contrast. In the Zhi et al. cohort, 115 of the 134 connections associated with HAMD in MDD-DIRECT (85.8%) show the same direction of association with HDRS total score. Associations with HAMA are concentrated in the somatomotor domain, which participates in 82 of the 246 connections, and are strikingly consistent in direction: coupling between subcortical and somatomotor networks increases with anxiety severity across SC-ET–SM (12 of 12 connections positive), SC-EH–SM (13 of 14), SC-BG–SM (9 of 9) and CB–SM (21 of 21). The strongest associations along this axis are between thalamus/hippocampus/amygdala and supplementary motor area (*r* = +0.109) and between left thalamus and frontal eye field (*r* = +0.106). This is the same subcortical-to-somatomotor axis identified in the diagnostic contrast, and it is the one place in our data where a diagnostic effect appears to scale with symptom burden. Consistent with this, and computed on the unthresholded statistics so as to be independent of any threshold, the HAMA effect map resembles the diagnostic effect map appreciably more than the HAMD map does (*r* = 0.352 against *r* = 0.184), suggesting that this axis may track anxiety rather than depression severity.

**Fig. 6.**
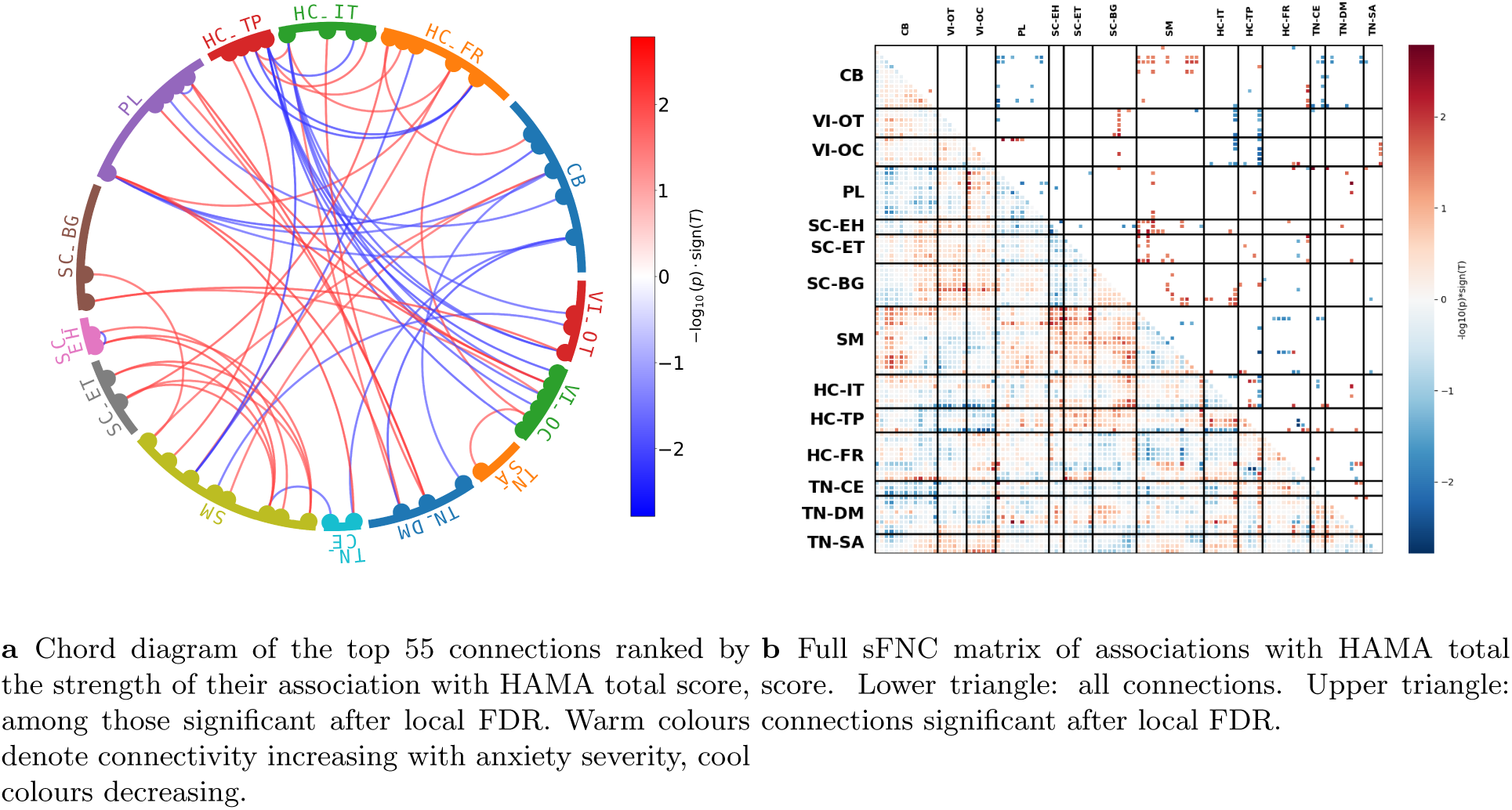
Associations between sFNC and HAMA total score in MDD-DIRECT. Patients with MDD and HAMA scores (*n ≈* 663). Each connection was regressed on HAMA total score, adjusting for age, sex and site, with local FDR correction (*q ≤* 0.05) across all 5,460 connections; effect size is the partial correlation *r*.

**Fig. 7.**
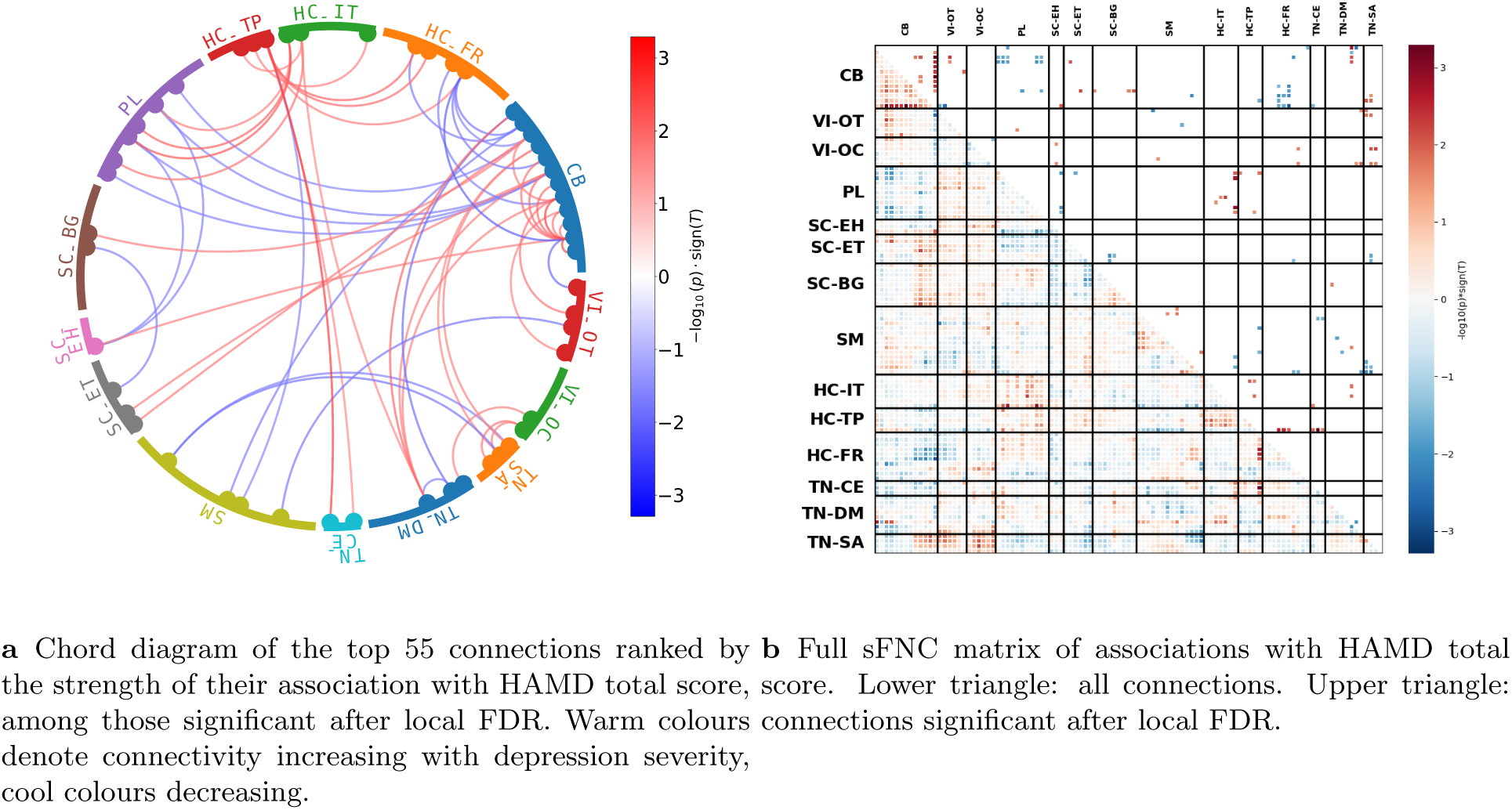
Associations between sFNC and HAMD total score in MDD-DIRECT. Patients with MDD and HAMD scores (*n ≈* 925). Each connection was regressed on HAMD total score, adjusting for age, sex and site, with local FDR correction (*q ≤* 0.05) across all 5,460 connections; effect size is the partial correlation *r*.

**Table 6.** Connections significantly associated with HAMD total score in MDD-DIRECT, by domain (local FDR *q ≤* 0.05; 134 connections in total). A connection is counted under each of its two domains. *N* ^+^ and *N ^−^*: connections whose coupling increases or decreases with severity. Effect size is the covariate-adjusted partial correlation *r*; Min *p* is uncorrected. Linear regression on HAMD total score adjusting for age, sex and site, in patients with MDD and HAMD scores (*n ≈* 925).

| Domain | Connections | *N*<sup>+</sup> | *N*<sup>−</sup> | Mean \|*r*\| | Max \|*r*\| | Min *p* |
|---|---|---|---|---|---|---|
| CB | 60 | 28 | 32 | 0.083 | 0.113 | 5.55 × 10<sup>−4</sup> |
| VI-OT | 12 | 8 | 4 | 0.080 | 0.096 | 3.50 × 10<sup>−3</sup> |
| VI-OC | 12 | 11 | 1 | 0.077 | 0.090 | 6.07 × 10<sup>−3</sup> |
| PL | 28 | 11 | 17 | 0.080 | 0.106 | 1.21 × 10<sup>−3</sup> |
| SC-EH | 6 | 3 | 3 | 0.077 | 0.085 | 1.00 × 10<sup>−2</sup> |
| SC-ET | 9 | 2 | 7 | 0.079 | 0.089 | 6.59 × 10<sup>−3</sup> |
| SC-BG | 7 | 4 | 3 | 0.077 | 0.086 | 8.88 × 10<sup>−3</sup> |
| SM | 25 | 6 | 19 | 0.077 | 0.091 | 5.69 × 10<sup>−3</sup> |
| HC-IT | 15 | 13 | 2 | 0.082 | 0.106 | 1.21 × 10<sup>−3</sup> |
| HC-TP | 15 | 12 | 3 | 0.083 | 0.114 | 5.15 × 10<sup>−4</sup> |
| HC-FR | 20 | 5 | 15 | 0.082 | 0.104 | 1.59 × 10<sup>−3</sup> |
| TN-CE | 7 | 4 | 3 | 0.080 | 0.114 | 5.15 × 10<sup>−4</sup> |
| TN-DM | 16 | 10 | 6 | 0.081 | 0.099 | 2.50 × 10<sup>−3</sup> |
| TN-SA | 17 | 10 | 7 | 0.079 | 0.091 | 5.69 × 10<sup>−3</sup> |

**Table 7.** Connections significantly associated with HAMA total score in MDD-DIRECT, by domain (local FDR *q ≤* 0.05; 246 connections in total). A connection is counted under each of its two domains. *N* ^+^ and *N ^−^*: connections whose coupling increases or decreases with severity. Effect size is the covariate-adjusted partial correlation *r*; Min *p* is uncorrected. Linear regression on HAMA total score adjusting for age, sex and site, in patients with MDD and HAMA scores (*n ≈* 663).

| Domain | Connections | *N*<sup>+</sup> | *N*<sup>−</sup> | Mean \|*r*\| | Max \|*r*\| | Min *p* |
|---|---|---|---|---|---|---|
| CB | 60 | 27 | 33 | 0.086 | 0.110 | 4.68 × 10<sup>−3</sup> |
| VI-OT | 20 | 8 | 12 | 0.089 | 0.107 | 5.79 × 10<sup>−3</sup> |
| VI-OC | 23 | 11 | 12 | 0.094 | 0.114 | 3.33 × 10<sup>−3</sup> |
| PL | 39 | 17 | 22 | 0.090 | 0.122 | 1.65 × 10<sup>−3</sup> |
| SC-EH | 22 | 16 | 6 | 0.088 | 0.109 | 5.09 × 10<sup>−3</sup> |
| SC-ET | 22 | 18 | 4 | 0.086 | 0.106 | 6.55 × 10<sup>−3</sup> |
| SC-BG | 34 | 29 | 5 | 0.086 | 0.107 | 5.79 × 10<sup>−3</sup> |
| SM | 82 | 63 | 19 | 0.087 | 0.116 | 2.73 × 10<sup>−3</sup> |
| HC-IT | 40 | 20 | 20 | 0.089 | 0.107 | 5.94 × 10<sup>−3</sup> |
| HC-TP | 41 | 19 | 22 | 0.090 | 0.119 | 2.11 × 10<sup>−3</sup> |
| HC-FR | 41 | 19 | 22 | 0.089 | 0.119 | 2.11 × 10<sup>−3</sup> |
| TN-CE | 20 | 6 | 14 | 0.087 | 0.122 | 1.65 × 10<sup>−3</sup> |
| TN-DM | 27 | 17 | 10 | 0.090 | 0.118 | 2.48 × 10<sup>−3</sup> |
| TN-SA | 9 | 6 | 3 | 0.084 | 0.096 | 1.34 × 10<sup>−2</sup> |

**Table 8.** Connections significantly associated with HAMD total score in MDD-DIRECT (local FDR *q ≤* 0.05), by pair of domains. Diagonal entries are within-domain connections. Linear regression on HAMD total score adjusting for age, sex and site, in patients with MDD and HAMD scores (*n ≈* 925).

| Domain | CB | VI-OT | VI-OC | PL | SC-EH | SC-ET | SC-BG | SM | HC-IT | HC-TP | HC-FR | TN-CE | TN-DM | TN-SA |
|---|---|---|---|---|---|---|---|---|---|---|---|---|---|---|
| CB | 13 | 5 | 0 | 13 | 1 | 2 | 3 | 1 | 0 | 1 | 12 | 1 | 5 | 3 |
| VI-OT | 5 | 0 | 0 | 1 | 0 | 0 | 0 | 2 | 0 | 0 | 0 | 0 | 1 | 3 |
| VI-OC | 0 | 0 | 1 | 0 | 2 | 0 | 0 | 1 | 0 | 0 | 2 | 0 | 2 | 4 |
| PL | 13 | 1 | 0 | 0 | 3 | 1 | 0 | 0 | 7 | 3 | 0 | 0 | 0 | 0 |
| SC-EH | 1 | 0 | 2 | 3 | 0 | 0 | 0 | 0 | 0 | 0 | 0 | 0 | 0 | 0 |
| SC-ET | 2 | 0 | 0 | 1 | 0 | 0 | 3 | 0 | 0 | 0 | 1 | 0 | 0 | 2 |
| SC-BG | 3 | 0 | 0 | 0 | 0 | 3 | 0 | 0 | 0 | 0 | 0 | 0 | 1 | 0 |
| SM | 1 | 2 | 1 | 0 | 0 | 0 | 0 | 4 | 2 | 2 | 2 | 2 | 4 | 5 |
| HC-IT | 0 | 0 | 0 | 7 | 0 | 0 | 0 | 2 | 0 | 3 | 0 | 1 | 2 | 0 |
| HC-TP | 1 | 0 | 0 | 3 | 0 | 0 | 0 | 2 | 3 | 0 | 3 | 3 | 0 | 0 |
| HC-FR | 12 | 0 | 2 | 0 | 0 | 1 | 0 | 2 | 0 | 3 | 0 | 0 | 0 | 0 |
| TN-CE | 1 | 0 | 0 | 0 | 0 | 0 | 0 | 2 | 1 | 3 | 0 | 0 | 0 | 0 |
| TN-DM | 5 | 1 | 2 | 0 | 0 | 0 | 1 | 4 | 2 | 0 | 0 | 0 | 1 | 0 |
| TN-SA | 3 | 3 | 4 | 0 | 0 | 2 | 0 | 5 | 0 | 0 | 0 | 0 | 0 | 0 |

**Table 9.** Connections significantly associated with HAMA total score in MDD-DIRECT (local FDR *q ≤* 0.05), by pair of domains. Diagonal entries are within-domain connections. Linear regression on HAMA total score adjusting for age, sex and site, in patients with MDD and HAMA scores (*n ≈* 663).

| Domain | CB | VI-OT | VI-OC | PL | SC-EH | SC-ET | SC-BG | SM | HC-IT | HC-TP | HC-FR | TN-CE | TN-DM | TN-SA |
|---|---|---|---|---|---|---|---|---|---|---|---|---|---|---|
| CB | 1 | 0 | 0 | 14 | 0 | 0 | 0 | 21 | 1 | 2 | 5 | 8 | 7 | 1 |
| VI-OT | 0 | 0 | 0 | 1 | 0 | 0 | 8 | 1 | 4 | 5 | 0 | 1 | 0 | 0 |
| VI-OC | 0 | 0 | 0 | 4 | 0 | 0 | 1 | 0 | 6 | 6 | 2 | 0 | 0 | 4 |
| PL | 14 | 1 | 4 | 4 | 1 | 1 | 1 | 4 | 0 | 1 | 2 | 1 | 4 | 1 |
| SC-EH | 0 | 0 | 0 | 1 | 1 | 1 | 2 | 14 | 1 | 0 | 1 | 0 | 1 | 0 |
| SC-ET | 0 | 0 | 0 | 1 | 1 | 0 | 0 | 12 | 0 | 2 | 5 | 0 | 1 | 0 |
| SC-BG | 0 | 8 | 1 | 1 | 2 | 0 | 0 | 9 | 8 | 2 | 2 | 0 | 0 | 1 |
| SM | 21 | 1 | 0 | 4 | 14 | 12 | 9 | 3 | 1 | 4 | 10 | 2 | 0 | 1 |
| HC-IT | 1 | 4 | 6 | 0 | 1 | 0 | 8 | 1 | 2 | 10 | 1 | 3 | 3 | 0 |
| HC-TP | 2 | 5 | 6 | 1 | 0 | 2 | 2 | 4 | 10 | 0 | 6 | 1 | 2 | 0 |
| HC-FR | 5 | 0 | 2 | 2 | 1 | 5 | 2 | 10 | 1 | 6 | 0 | 2 | 5 | 0 |
| TN-CE | 8 | 1 | 0 | 1 | 0 | 0 | 0 | 2 | 3 | 1 | 2 | 0 | 2 | 0 |
| TN-DM | 7 | 0 | 0 | 4 | 1 | 1 | 0 | 0 | 3 | 2 | 5 | 2 | 1 | 1 |
| TN-SA | 1 | 0 | 4 | 1 | 0 | 0 | 1 | 1 | 0 | 0 | 0 | 0 | 1 | 0 |

Associations with HAMD are instead concentrated in the cerebellum, which participates in 60 of the 134 connections. The strongest individual associations are between bilateral temporoparietal junction and left inferior parietal/dorsolateral prefrontal cortex (*r* = +0.114) and between dorsomedial cerebellum and inferior left ventrolateral prefrontal cortex (*r* = *−*0.104).

### Spectra

Spectral associations with symptom severity followed the same pattern as connectivity: present but sparse and weak relative to the diagnostic contrast. 347 of 13,545 bins (2.6%) were associated with HAMA total

score, 217 positively and 130 negatively, and 338 (2.5%) with HAMD total score, 164 positively and 174 negatively (Fig. 8; Tables 10–13). Maximum partial correlations were 0.144 for HAMA and 0.136 for HAMD. In the Zhi et al. cohort, 233 of the 338 bins associated with HAMD (68.9%) show the same direction of association with HDRS total score.

**Fig. 8.**
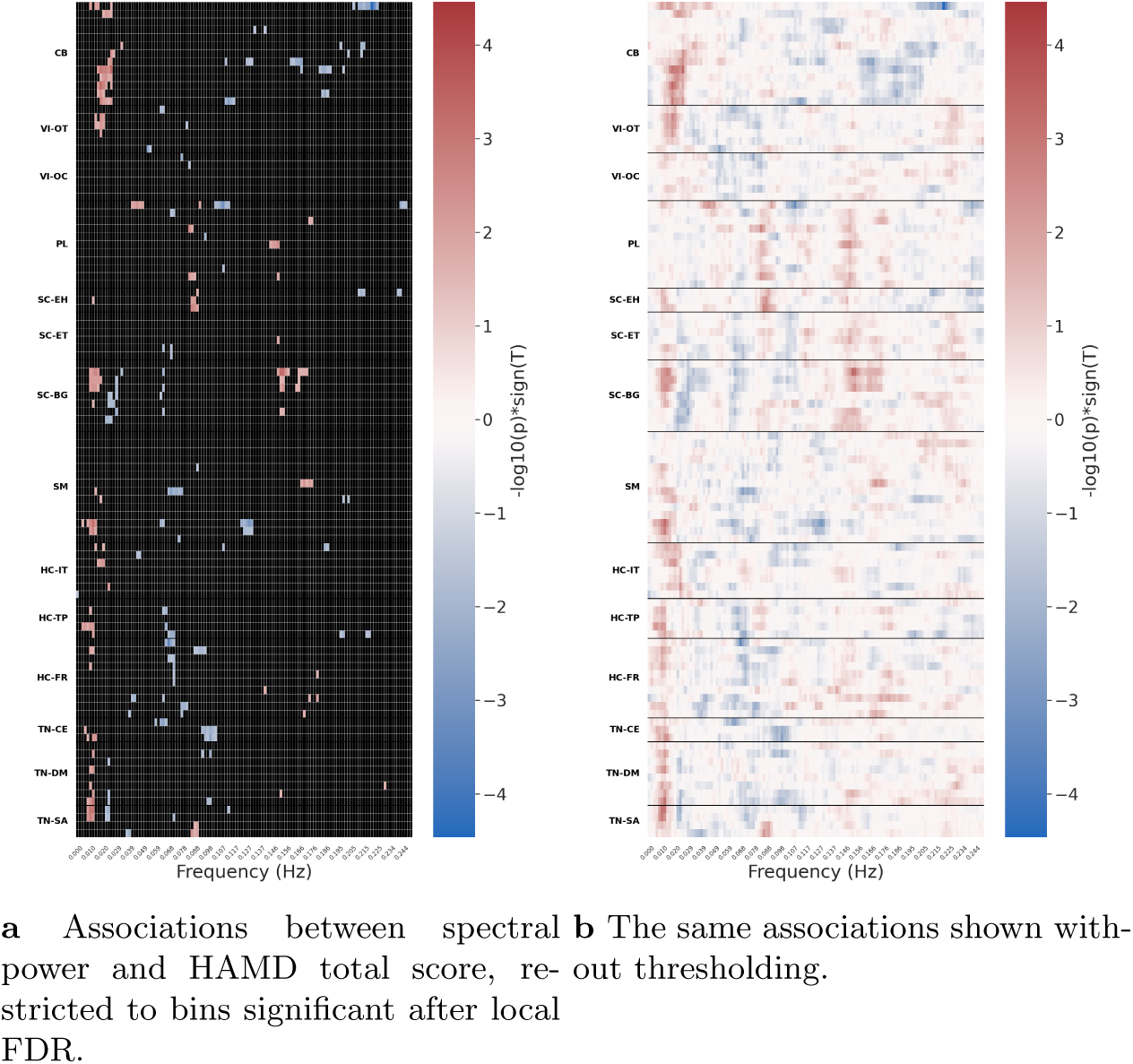
Associations between spectral power and HAMD total score in MDD-DIRECT. Patients with MDD and HAMD scores (*n ≈* 925). Power in each frequency bin of each network was regressed on HAMD total score, adjusting for age, sex and site, with local FDR correction (*q ≤* 0.05) across all 13,545 network–frequency tests; effect size is the partial correlation *r*.

**Table 10.** Spectral bins significantly associated with HAMA total score in MDD-DIRECT, by frequency range (local FDR *q ≤* 0.05; 347 bins in total). *N* ^+^ and *N ^−^*: bins whose power increases or decreases with severity. Effect size is the covariate-adjusted partial correlation *r*; Min *p* is uncorrected. Linear regression on HAMA total score adjusting for age, sex and site, in patients with MDD and HAMA scores (*n ≈* 663).

| Frequency range | Bins | *N*<sup>+</sup> | *N*<sup>−</sup> | Mean \|*r*\| | Max \|*r*\| | Min *p* |
|---|---|---|---|---|---|---|
| 0.00–0.05 Hz | 89 | 6 | 83 | 0.102 | 0.140 | 2.94 × 10<sup>−4</sup> |
| 0.05–0.10 Hz | 44 | 25 | 19 | 0.100 | 0.144 | 1.97 × 10<sup>−4</sup> |
| 0.10–0.15 Hz | 123 | 123 | 0 | 0.100 | 0.140 | 3.18 × 10<sup>−4</sup> |
| 0.15–0.20 Hz | 31 | 18 | 13 | 0.094 | 0.118 | 2.49 × 10<sup>−3</sup> |
| 0.20–0.25 Hz | 60 | 45 | 15 | 0.097 | 0.125 | 1.31 × 10<sup>−3</sup> |

**Table 11.** Spectral bins significantly associated with HAMA total score in MDD-DIRECT, by domain (local FDR *q ≤* 0.05; 347 bins in total). *N* ^+^ and *N ^−^*: bins whose power increases or decreases with severity. Effect size is the covariate-adjusted partial correlation *r*; Min *p* is uncorrected. Linear regression on HAMA total score adjusting for age, sex and site, in patients with MDD and HAMA scores (*n ≈* 663).

| Domain | Bins | *N*<sup>+</sup> | *N*<sup>−</sup> | Mean \|*r*\| | Max \|*r*\| | Min *p* |
|---|---|---|---|---|---|---|
| CB | 17 | 8 | 9 | 0.092 | 0.100 | 9.95 × 10<sup>−3</sup> |
| VI-OT | 20 | 8 | 12 | 0.101 | 0.144 | 1.97 × 10<sup>−4</sup> |
| VI-OC | 21 | 15 | 6 | 0.102 | 0.118 | 2.46 × 10<sup>−3</sup> |
| PL | 38 | 23 | 15 | 0.098 | 0.113 | 3.53 × 10<sup>−3</sup> |
| SC-EH | 4 | 1 | 3 | 0.102 | 0.112 | 3.92 × 10<sup>−3</sup> |
| SC-ET | 16 | 10 | 6 | 0.092 | 0.106 | 6.60 × 10<sup>−3</sup> |
| SC-BG | 31 | 28 | 3 | 0.101 | 0.140 | 3.18 × 10<sup>−4</sup> |
| SM | 36 | 27 | 9 | 0.098 | 0.130 | 8.36 × 10<sup>−4</sup> |
| HC-IT | 60 | 33 | 27 | 0.100 | 0.131 | 7.38 × 10<sup>−4</sup> |
| HC-TP | 3 | 3 | 0 | 0.097 | 0.104 | 7.45 × 10<sup>−3</sup> |
| HC-FR | 25 | 11 | 14 | 0.100 | 0.140 | 3.10 × 10<sup>−4</sup> |
| TN-CE | 16 | 15 | 1 | 0.100 | 0.121 | 1.83 × 10<sup>−3</sup> |
| TN-DM | 29 | 21 | 8 | 0.098 | 0.129 | 9.06 × 10<sup>−4</sup> |
| TN-SA | 31 | 14 | 17 | 0.107 | 0.140 | 2.94 × 10<sup>−4</sup> |

**Table 12.** Spectral bins significantly associated with HAMD total score in MDD-DIRECT, by frequency range (local FDR *q ≤* 0.05; 338 bins in total). *N* ^+^ and *N ^−^*: bins whose power increases or decreases with severity. Effect size is the covariate-adjusted partial correlation *r*; Min *p* is uncorrected. Linear regression on HAMD total score adjusting for age, sex and site, in patients with MDD and HAMD scores (*n ≈* 925).

| Frequency range | Bins | *N*<sup>+</sup> | *N*<sup>−</sup> | Mean \|*r*\| | Max \|*r*\| | Min *p* |
|---|---|---|---|---|---|---|
| 0.00–0.05 Hz | 137 | 109 | 28 | 0.086 | 0.115 | 4.54 × 10<sup>−4</sup> |
| 0.05–0.10 Hz | 86 | 17 | 69 | 0.083 | 0.118 | 3.37 × 10<sup>−4</sup> |
| 0.10–0.15 Hz | 36 | 4 | 32 | 0.084 | 0.113 | 5.75 × 10<sup>−4</sup> |
| 0.15–0.20 Hz | 54 | 33 | 21 | 0.081 | 0.113 | 5.79 × 10<sup>−4</sup> |
| 0.20–0.25 Hz | 25 | 1 | 24 | 0.085 | 0.136 | 3.45 × 10<sup>−5</sup> |

**Table 13.** Spectral bins significantly associated with HAMD total score in MDD-DIRECT, by domain (local FDR *q ≤* 0.05; 338 bins in total). *N* ^+^ and *N ^−^*: bins whose power increases or decreases with severity. Effect size is the covariate-adjusted partial correlation *r*; Min *p* is uncorrected. Linear regression on HAMD total score adjusting for age, sex and site, in patients with MDD and HAMD scores (*n ≈* 925).

| Domain | Bins | *N*<sup>+</sup> | *N*<sup>−</sup> | Mean \|*r*\| | Max \|*r*\| | Min *p* |
|---|---|---|---|---|---|---|
| CB | 76 | 37 | 39 | 0.087 | 0.136 | 3.45 × 10<sup>−5</sup> |
| VI-OT | 13 | 8 | 5 | 0.084 | 0.099 | 2.61 × 10<sup>−3</sup> |
| VI-OC | 2 | 0 | 2 | 0.083 | 0.090 | 6.17 × 10<sup>−3</sup> |
| PL | 31 | 18 | 13 | 0.085 | 0.113 | 5.75 × 10<sup>−4</sup> |
| SC-EH | 12 | 7 | 5 | 0.083 | 0.099 | 2.58 × 10<sup>−3</sup> |
| SC-ET | 4 | 1 | 3 | 0.078 | 0.080 | 1.50 × 10<sup>−2</sup> |
| SC-BG | 49 | 32 | 17 | 0.083 | 0.113 | 5.79 × 10<sup>−4</sup> |
| SM | 36 | 15 | 21 | 0.085 | 0.112 | 6.31 × 10<sup>−4</sup> |
| HC-IT | 12 | 6 | 6 | 0.081 | 0.093 | 4.83 × 10<sup>−3</sup> |
| HC-TP | 17 | 7 | 10 | 0.083 | 0.090 | 6.06 × 10<sup>−3</sup> |
| HC-FR | 31 | 8 | 23 | 0.081 | 0.118 | 3.37 × 10<sup>−4</sup> |
| TN-CE | 19 | 4 | 15 | 0.083 | 0.099 | 2.50 × 10<sup>−3</sup> |
| TN-DM | 17 | 10 | 7 | 0.084 | 0.110 | 8.21 × 10<sup>−4</sup> |
| TN-SA | 19 | 11 | 8 | 0.086 | 0.109 | 9.26 × 10<sup>−4</sup> |

The two scales again implicate different networks. HAMA associations are concentrated in insular net-works: right insula, ventral anterior insula and medial ventral anterior insula together account for 38 bins, predominantly showing increased power with increasing anxiety. These associations appear alongside alterations in the right extrastriate cortex (13 bins), left medial temporal pole (17) and left caudate (7, all positive). HAMD associations are concentrated in the cerebellum (76 bins across the anterior, anterior dorsomedial, medial and dorsomedial cerebellar networks) and lateral temporal pole (15), echoing the cerebellar concentration of the HAMD connectivity associations. Within each network, however, the associated bins are distributed across much of the spectrum with mixed sign, in contrast to the narrow, direction-consistent bands seen in the diagnostic analysis.

### Spatial Maps

The same effect-size criterion was applied to the severity analyses, using the partial correlation in place of Cohen’s *d*. For HAMA, 89,642 voxels reach *|r| ≥* 0.10 and 1,189 reach *|r| ≥* 0.15, but only **six** reach *|r| ≥* 0.20 and none reach 0.25; the largest partial correlation anywhere in the brain is 0.231. For HAMD the corresponding counts are 20,753, 83 and **zero**, with a maximum partial correlation of 0.193. Restricted to the template extents (*|Z| >* 2), 63,841 voxel–network tests survive local FDR for HAMA and 112,812 for HAMD, none with *|r| ≥* 0.2 (Figs. 9 and 10; Tables 14 and 15). Spatial-map associations with HDRS total score in the Zhi et al. cohort are shown in Supplementary Fig. 5.

**Fig. 9.**
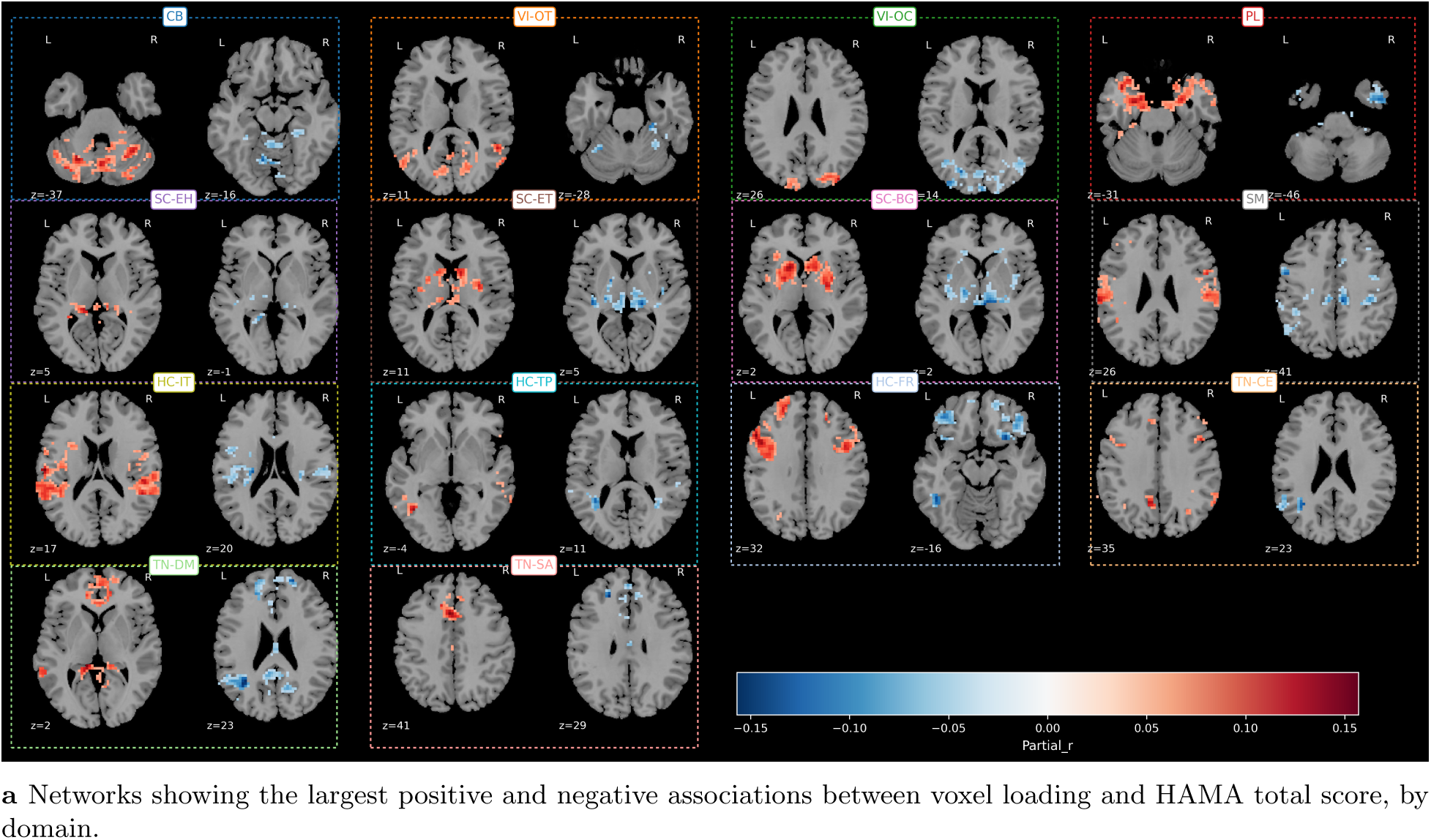
Associations between spatial map intensity and HAMA total score in MDD-DIRECT. Patients with MDD and HAMA scores (*n ≈* 663). Voxel loadings were regressed on HAMA total score, adjusting for age, sex and site. Voxels are restricted to each network’s NeuroMark 2.2 template extent (*|Z| >* 2) and shown where significant after local FDR (*q ≤* 0.05); colour shows the covariate-adjusted partial correlation *r*.

**Fig. 10.**
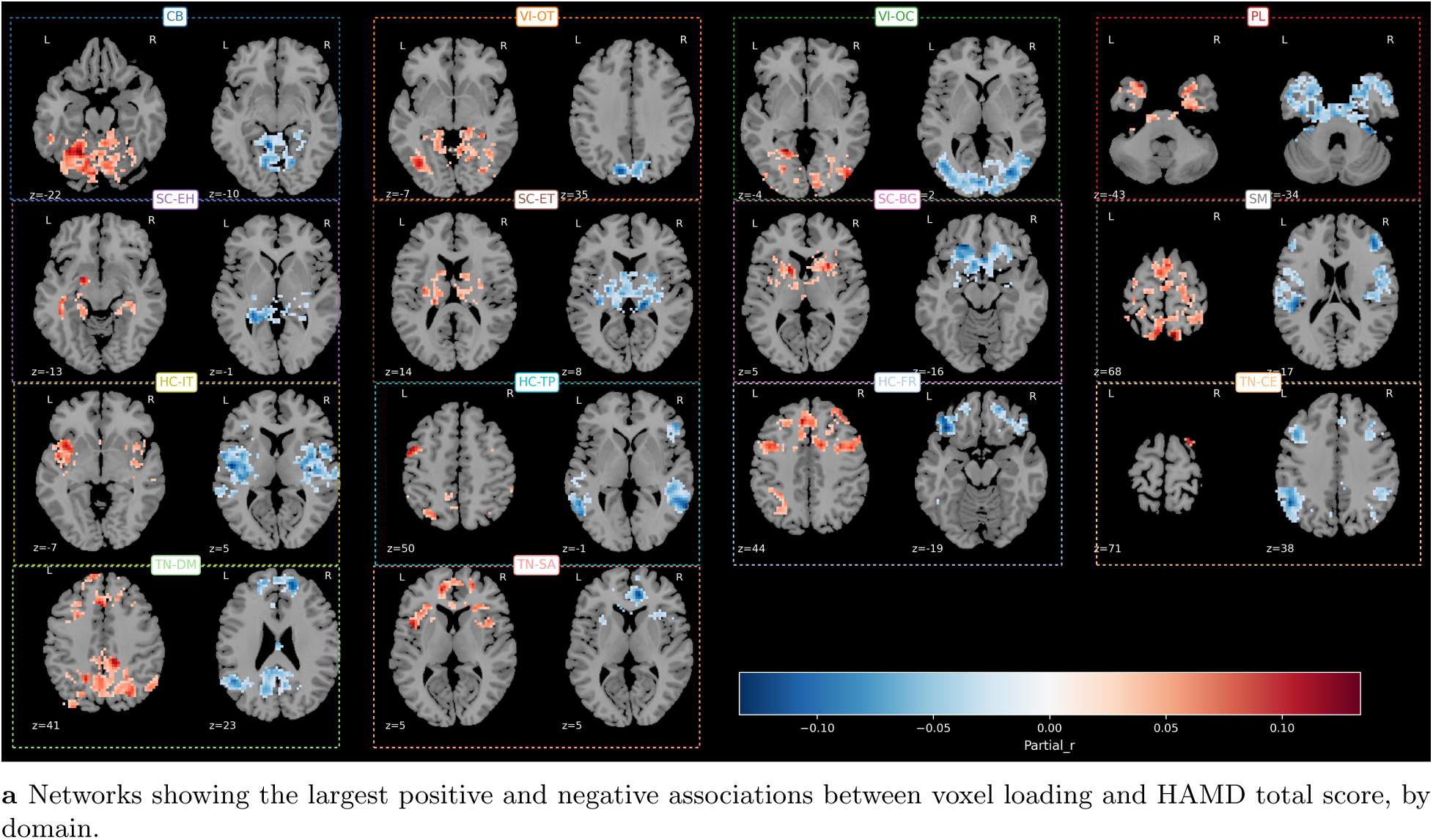
Associations between spatial map intensity and HAMD total score in MDD-DIRECT. Patients with MDD and HAMD scores (*n ≈* 925). Voxel loadings were regressed on HAMD total score, adjusting for age, sex and site. Voxels are restricted to each network’s NeuroMark 2.2 template extent (*|Z| >* 2) and shown where significant after local FDR (*q ≤* 0.05); colour shows the covariate-adjusted partial correlation *r*.

**Table 14.** Number of distinct voxels significant after local FDR (*q ≤* 0.05) for the HAMD total score, by domain, with each network restricted to its NeuroMark 2.2 template extent. (*|Z| >* 2). Voxelwise regression on HAMD total score adjusting for age, sex and site, in patients with MDD and HAMD scores (*n ≈* 925).

| Domain | Significant voxels |
|---|---|
| CB | 6761 |
| VI-OT | 4782 |
| VI-OC | 5812 |
| PL | 5118 |
| SC-EH | 1365 |
| SC-ET | 2710 |
| SC-BG | 4103 |
| SM | 11436 |
| HC-IT | 4886 |
| HC-TP | 4220 |
| HC-FR | 8248 |
| TN-CE | 3558 |
| TN-DM | 8222 |
| TN-SA | 3033 |

**Table 15.**
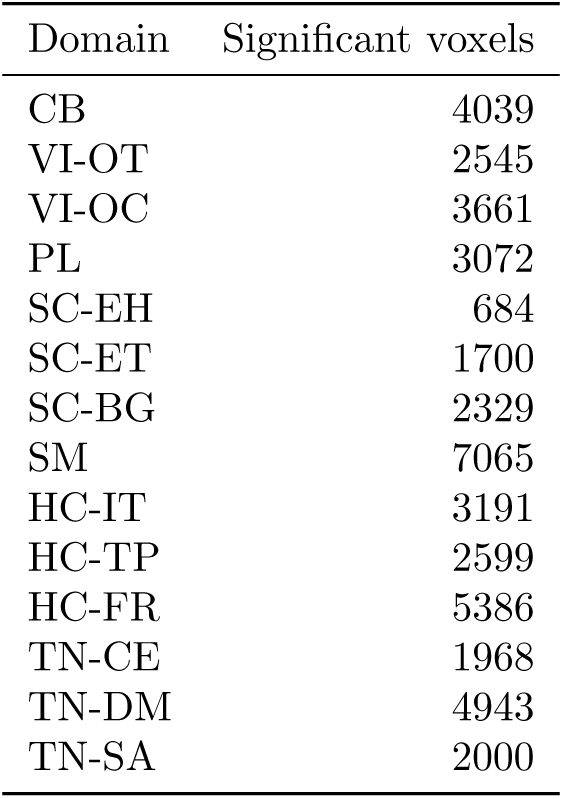
Number of distinct voxels significant after local FDR (*q ≤* 0.05) for the HAMA total score, by domain, with each network restricted to its NeuroMark 2.2 template extent. (*|Z| >* 2). Voxelwise regression on HAMA total score adjusting for age, sex and site, in patients with MDD and HAMA scores (*n ≈* 663).

| Domain | Significant voxels |
|---|---|
| CB | 4039 |
| VI-OT | 2545 |
| VI-OC | 3661 |
| PL | 3072 |
| SC-EH | 684 |
| SC-ET | 1700 |
| SC-BG | 2329 |
| SM | 7065 |
| HC-IT | 3191 |
| HC-TP | 2599 |
| HC-FR | 5386 |
| TN-CE | 1968 |
| TN-DM | 4943 |
| TN-SA | 2000 |

The local-FDR threshold does not constrain these results: from *|r| ≥* 0.10 downward the counts are identical for every threshold between *q ≤* 0.05 and *q ≤* 0.001, so what follows concerns the magnitude of the associations rather than the correction applied. At the lower end of the range the associations are widespread, with 89,642 voxels for HAMA (1.19% of tests) and 20,753 for HAMD (0.28%), and all 105 networks represented for both scales.

Weak as they are, these associations are not anatomically arbitrary, and for each scale they fall in the systems that scale implicates in the other two derivatives. The HAMA associations at *|r| ≥* 0.10 are most concentrated in the cerebellar (1.27% of voxels), insular (1.24%) and occipitotemporal visual (1.24%) domains, with the largest proportions in bilateral cerebellum (1.59%), paracentral lobule (1.53%), anterior cerebellum (1.53%), medial ventral anterior insula (1.50%) and right inferior parietal lobe/dorsolateral prefrontal cortex (1.49%). The paracentral and insular involvement parallels the somatomotor and insular emphasis of the HAMA connectivity and spectral results, and the right inferior parietal/dorsolateral prefrontal network is the same one that supplies five of the six voxels surviving at *|r| ≥* 0.20. The HAMD associations are most concentrated in the cerebellum (0.31% of voxels, with six of the twelve most affected networks cerebellar) and the salience domain (0.31%), and several of the cerebellar networks are strongly direction-dominant: anterior cerebellum (187 of 280 voxels positive), ventromedial cerebellum (175 of 272) and right extrastriate cortex (178 of 272). This cerebellar concentration recapitulates the HAMD connectivity result, in which the cerebellum participated in 60 of 134 connections, and the HAMD spectral result, in which cerebellar networks contributed the largest group of associated bins. Three methodologically distinct derivatives therefore converge on the cerebellum for depression severity, which is a considerably stronger basis for that observation than any one of them alone.

## Discussion

In this work, we have performed a comprehensive study of a large-scale resting-state fMRI dataset, MDDDIRECT. In contrast to previous studies which focused on targeted regions or specific analysis frameworks, our study is full-brain and analyzes the changes in MDD using multiple derivatives: static functional network connectivity (sFNC), time-series spectra, and ICA spatial maps. We are primarily interested in assessing the variations in these derivatives between MDD patients and controls. As diagnosis can be notoriously fickle, we are also interested in studying the associations of these derivatives with symptom severity. We also assess the consistency of these findings in the smaller cohort from Zhi et al. [19]. In the following sections, we discuss each sFNC, spectra and spatial map differences individually, and then discuss potential interactions between these findings. Finally, we discuss the limitations and directions for future studies which can address these limitations and expand on the analysis we perform here.

### MDD and Functional Network Connectivity

Large, singular studies of MDD, such as the one we present in this work, are rare. In smaller cohorts and meta-analyses, however, the effects of major depressive disorder on functional connectivity have been widely studied. Despite small sample sizes, studies targeting specific regions identified consistent effects in MDD, such as increased connectivity within the default mode, increased default-mode/salience connectivity, and decreased default-mode/central-executive connectivity[36]. Analyses outside the default mode and saliency networks indicate wide-spread functional network dysfunction[17] as well as an overall story of “abnormal connectivity” in which different regions are found to more or less connected in MDD than in controls.

In contrast to these smaller studies, the large (*N* = 2428) MDD-Rest study identified decreased connectivity within the default mode [37], driven by individuals with recurrent depression and undergoing medical treatment. This inconsistency between the findings in large and small studies suggests that more evidence is needed, with more attention given to potential confounding factors that can complicate particular findings.

Across 5460 network pairs we find differences that are widespread but small: 11.9% of connections differ, none exceeding *|d|* = 0.41, and with a mixture of increases and decreases rather than a uniform direction.

This is consistent with the “abnormal connectivity” characterization of the meta-analytic literature[17], and it also suggests why regionally targeted studies in small samples have been difficult to reconcile: at these effect sizes, a study powered to detect *|d|* = 0.5 in a handful of *a priori* regions will recover a different and largely arbitrary subset of the true effect map in each sample.

The structure we observe is more specific than diffuse dysconnectivity. Four blocks account for a large share of the significant connections and each is nearly uniform in direction. The most prominent is a reduction in cerebello-subcortical coupling, affecting essentially every significant cerebellum–basal ganglia and cerebellum–thalamus connection and driven most strongly by connections between cerebellar networks and the caudate. This is the cerebellar limb of the cortico-striato-thalamo-cerebellar loop which has been consistently identified within a growing literature on cerebellar contributions to affective disorder [20, 21, 23]. In the opposite direction, coupling between thalamic, hippocampal-amygdalar and striatal networks and somatomotor cortex is almost uniformly increased. A third block comprises reduced coupling between visual networks and posterior temporal, temporoparietal and precuneus networks, and contains both the largest and the most statistically reliable effects in our analysis. A fourth comprises increased coupling between hippocampal, parahippocampal and amygdalar networks and inferior parietal and postcentral cortex, the most directionally consistent gain of coupling in the study and the finding most directly connected to affective and salience accounts of depression.

In contrast to their prevalence in the literature, the three triple-network domains, central executive, default mode and salience, are the least affected of all domains, with 13 of 309, 44 of 804 and 39 of 410 connections significant respectively, against a whole-brain rate of 11.9%. We note that our findings are not inconsistent with [37], in whose large sample the default-mode effect was small and confined to recurrent, medicated patients; effects that are small and distributed over a small number of network pairs will contribute few significant connections in a whole-brain analysis even when they are real. Our findings suggest, however, that at the whole-brain scale the connectivity signature of MDD is dominated by thalamic, cerebellar and sensorimotor systems rather than by the associative networks that have received most attention.

### MDD and Spectra

Frequency-domain properties of the BOLD signal have received considerably less attention in MDD than connectivity, and where they have been studied it has usually been through summary measures such as ALFF or fALFF computed within a fixed low-frequency band [20, 21, 22, 24], or through frequency-resolved connectivity [31]. Analyzing the full spectrum bin by bin, we find that the dominant effect of MDD is not a change in the amount of low-frequency power but a change in how power is distributed across frequencies. Power is reduced at low frequencies and increased at high frequencies, so that the group difference is best described as a flattening of the BOLD power spectrum rather than a gain or loss of signal amplitude.

The spectral effect is concentrated in the extended thalamus, the same domain that shows the highest proportion of altered connections. Within thalamic networks the frequency dependence is explicit, with reductions confined to frequencies below roughly 0.03 Hz and increases appearing above 0.15 Hz. The convergence of two methodologically independent derivatives on the same anatomical target is the strongest evidence in this study for thalamic involvement in MDD. The thalamus is well positioned to produce both effects at once: as the principal relay between subcortical and cortical systems, a change in its intrinsic oscillatory regime would be expected to manifest both as altered spectral content and as altered coupling with its cortical targets, which is what we observe. Beyond the thalamus, the insula and cerebellum show reduced power near 0.10 Hz, and both are also prominent in the connectivity results.

### MDD and Spatial Variation of Functional Networks

The spatial maps estimated by NeuroMark ICA describe, for each network, how strongly each voxel contributes to that network’s time course. Group differences in these maps therefore index a property distinct from connectivity and spectra: not how networks interact or oscillate, but how they are spatially constituted. Our findings suggest that cerebellar, diencephalic and basal-ganglia networks gain voxel loading in MDD, while somatomotor and right parietal association networks lose it, and this directionality is highly consistent within networks: 50 of the 83 networks with an appreciable number of surviving voxels depart significantly from directional balance, and several are almost perfectly unidirectional, including bilateral basal ganglia, vermis, anterior diencephalon and superior sensorimotor cortex. Interpreted at face value, a gain of loading means that the voxels concerned contribute more strongly to that network’s time course in patients, and a loss that they contribute less; the pattern is therefore one in which subcortical and cerebellar systems become more spatially dominant while sensorimotor and parietal association systems become less so.

Two aspects of these differences need to be separated, because they point to different systems and a single threshold reports only one of them. Considered by extent, the differences are pervasive: at a minimum effect size of 0.1 every one of the 105 networks contains affected voxels, and on average 6.9% of each network’s extent differs between groups. Considered by magnitude, they are focal and cerebellar. The two orderings do not agree. By extent the default mode is the most affected domain, followed by the visual and thalamic domains with the salience domain close behind; by magnitude the cerebellum dominates, supplying more than a quarter of the voxels surviving at 0.2 and nearly half of those surviving at 0.25. The default mode and salience networks are therefore altered across a large fraction of their extent but weakly at any given voxel, while the cerebellum is altered strongly in a smaller number of voxels. Both are real, and describing only the second understates the involvement of precisely those systems the depression literature has most often implicated.

The gradient in directional consistency supports this two-component reading. Within-network agreement in the sign of the effect rises steadily with the effect-size floor, from 0.53 at 0.1 through 0.58 at 0.15 to 0.71 at 0.2, which is what one expects if focal and coherent changes sit within a broader, less organized background rather than if a single process operated uniformly across the brain. That the diffuse component is not simply noise is indicated by 69 of the 105 networks departing significantly from directional balance even at the lowest floor. The ordering shifts once more when the statistical rather than the effect-size threshold is tightened, with the extended thalamus becoming the most affected domain — the same domain that leads the connectivity and spectral analyses, and further evidence that thalamic involvement is the most statistically reliable feature of these data even where it is not the largest in magnitude.

This is worth setting alongside the connectivity findings, with which it partly agrees and partly does not. The cerebellum, diencephalon and basal ganglia are prominent in both derivatives, as is somatomotor cortex, so the same systems are implicated. In the connectivity analysis the cerebellum *decouples* from the subcortex while the subcortex *over-couples* to somatomotor cortex. In the spatial maps, functional networks in the cerebellum and basal ganglia both expand. A network can gain spatial extent while losing coupling to its neighbours, and the co-occurrence of the two in the cerebello-subcortical system is a specific, testable observation rather than a redundancy between derivatives. The triple-network domains, which are the least affected in connectivity, are also not prominent here once effect size is accounted for — although we note that they are more prominent at lower effect-size thresholds, and the appropriate reading is that their spatial-map differences are widespread but individually small.

The spatial-map findings are, however, the least consistent of our results across cohorts. Their direction is preserved, with 77% of voxels at the primary criterion agreeing in sign, but the unthresholded effect maps correlate only weakly (*r* = 0.15), and the smaller cohort’s own analysis does not recover the coherent within-network gains and losses seen in MDD-DIRECT (Supplementary Note 2). Two explanations are plausible and we cannot presently distinguish them. The first is statistical: standardized effect sizes are inflated at *N* = 519, and the voxelwise spatial-map test is the least favourable of our three derivatives in this respect, because it has by far the largest number of tests and the smallest per-test effect. The second is methodological: spatial maps are the derivative most sensitive to differences in normalization and smoothing between the two independently preprocessed cohorts, since they are estimated in voxel space rather than from summary time-series properties. Either way, the spatial-map results should be regarded as established in MDD-DIRECT and awaiting independent confirmation, and we weight them accordingly below.

### Symptom Severity

Associations with symptom severity were markedly sparser and weaker than the differences between patients and controls, across all three derivatives, despite being measured in the same participants with the same pipeline. Two readings are available and they are not mutually exclusive. The first is statistical: severity scores were available for only a subset of the cohort, so these analyses rest on roughly 663 and 925 participants against approximately 1,930 for the diagnostic contrast, and the largest severity effects we observe (partial correlations of 0.12 for connectivity and 0.14 for spectra) are small enough that a substantial reduction in sample size would be expected to attenuate them considerably.

The spatial maps qualify this picture in one respect. Although no voxel reaches a moderate effect size for either scale, the weak associations that are present are anatomically organized rather than scattered, and they fall in the systems each scale implicates elsewhere: insular and paracentral networks for HAMA, and the cerebellum for HAMD, where six of the twelve most affected networks are cerebellar and several are strongly direction-dominant. For depression severity in particular, connectivity, spectra and spatial composition all point to the cerebellum. Three derivatives computed from the same data are not independent evidence in the statistical sense, but they are sensitive to different properties of it, and their agreement carries more weight than any single weak result.

Connectivity is a partial exception, and an informative one. The HAMA associations reproduce the subcortical-to-somatomotor axis seen in the diagnostic contrast, with the same direction and the same participating domains, and the HAMA effect map resembles the diagnostic map appreciably more than the HAMD map does (*r* = 0.352 against *r* = 0.184). This suggests that one component of the diagnostic connectivity signature does scale with symptom burden, and that it does so with anxiety rather than depression severity. The HAMD associations instead implicate a distinct cerebello-prefrontal circuit, with coupling between cerebellum and ventrolateral prefrontal cortex decreasing and coupling within the cerebellum increasing as depression severity rises. We regard this dissociation as provisional, since the two scales were administered to overlapping but non-identical subsets and their effect maps agree less well than two clinically correlated instruments should. A parallel observation arises in the spatial maps, where five of the six voxels associated with HAMA fall in the right inferior parietal/dorsolateral prefrontal network and show reduced loading with increasing anxiety — the same network and direction that is strongly direction-dominant in the diagnostic contrast. These derivatives thus independently suggest that anxiety severity tracks a component of the diagnostic effect, and both point to systems that are already altered between groups.

### Cross-Cohort Consistency

We have treated the Zhi et al. cohort as an assessment of cross-cohort consistency. At the effect sizes MDD produces in these derivatives, a cohort of 519 participants has at best moderate power to detect any individual effect, so the number of findings significant in both cohorts is determined largely by the smaller cohort’s power. The direction of effect does not have this limitation, since an effect that is real and shared should carry the same sign in both cohorts whether or not it reaches significance in the smaller one.

On this criterion the connectivity findings are highly consistent: 95% of the connections significant in MDD-DIRECT, and 96% of those with at least a minor effect, have the same direction in Zhi et al. The spectral findings agree in direction at 78% of significant bins, with agreement concentrated below 0.15 Hz and in the thalamic, cerebellar and default-mode networks, and the spatial-map findings agree at 77% of voxels at the primary criterion. For connectivity and spatial maps, agreement rises with effect size, as expected if the larger effects are the more reliably estimated.

The cerebello-subcortical connectivity block is the densest finding in MDD-DIRECT, yet contributes no connection that is significant in both cohorts. Its direction, however, agrees in 75 of 81 connections, so the absence of overlap is better explained by the power of the smaller cohort than by inconsistency between cohorts.

### Limitations and Future Work

We would like to acknowledge some of the limitations of the current study, which highlight several important directions for future studies.

The most significant methodological limitation concerns preprocessing. The two cohorts were preprocessed independently according to their own study protocols, and although we matched smoothing parameters and normalization targets, the pipelines differ in the software used for several steps. Because small pipeline differences are known to propagate into derived measures, some portion of the cross-study discrepancy we report may be methodological rather than biological. Future multi-site efforts should adopt uniform preprocessing or, preferably, release raw anonymized data so that a common pipeline can be applied retrospectively.

Our analysis is also unimodal. Structural and diffusion measures are known to be altered in several of the regions we implicate, particularly the hippocampus and cerebellum, and functional data alone cannot determine whether the differences we observe reflect functional reorganization or the functional consequences of structural change. This caveat bears most directly on the spatial-map results, where a change in the estimated spatial extent of a network is precisely the kind of effect that regional gray matter differences could produce. A multimodal analysis incorporating gray matter volume is the natural next step.

The cohorts studied here are geographically and ethnically limited, comprising East Asian participants recruited in China. Whether the effect map we describe generalizes to other populations is an open question, and one that bears directly on the biomarker applications discussed above.

Finally, the symptom-severity analyses were limited to the subset of participants for whom scores were available, reducing the effective sample by roughly half relative to the diagnostic contrast, and the two severity scales were administered to overlapping but non-identical subsets. The severity results should be understood in that light.

Several future directions follow directly. The most immediate is a longitudinal, within-subject design capable of distinguishing trait from state effects, which the cross-sectional design used here cannot do. A multimodal extension incorporating structural and diffusion measures would address the interpretive ambiguity noted above. Replication and extension in cohorts from other geographic regions would establish generality. A multi-derivative predictive framework, combining connectivity and spectral features rather than treating them separately, is a natural use of the fact that the two derivatives converge anatomically while carrying partially independent information. Incorporating treatment response and medication status, extending to interventional contexts such as TMS, collecting broader clinical and behavioral measures to rule out confounds, and normative modeling of individual deviation from population trajectories are all valuable further directions.

## Conclusion

We have presented a whole-brain, multi-derivative analysis of major depressive disorder in a large multisite cohort, using multi-scale template-driven ICA to obtain 105 functional networks and examining their connectivity, spectral content and spatial composition within a single statistical framework. Differences between patients and controls were widespread but small in effect. We do identify some structure in our findings: cerebello-subcortical coupling was reduced while subcortical-somatomotor coupling was increased, visual networks were decoupled from posterior temporal and precuneus networks, and the spectral content of thalamic networks shifted from lower to higher frequencies. The extended thalamus emerged as the point of convergence between the connectivity and spectral results, and is the single most strongly implicated system in our data. Differences in network spatial composition were pervasive, present in every network examined, and separated into a widespread component most extensive in the default mode and salience networks and a focal, directionally coherent component in which cerebellar, diencephalic and basal-ganglia networks gained voxel loading while somatomotor and right parietal association networks lost it. Associations with symptom severity were sparser and weaker; the clearest, a subcortical-to-somatomotor connectivity axis that tracked anxiety severity, is a natural target for longitudinal follow-up. Finally, in a smaller independent cohort analyzed under an identical procedure, the direction of effect agreed for 95% of the significant connections, 78% of the significant spectral bins and 77% of the largest spatial-map effects, indicating that the characterization obtained in the large cohort is consistent across cohorts even where the smaller cohort lacks the power to detect the effects individually.

## Methods

First, data-driven experiments using a Multi-Scale NeuroMark Independent Component Analysis (NM-ICA) on the large MDD-DIRECT data set enable us to identify a robust, replicable set of functional networks in the brain. We study these functional networks to determine how MDD can manifest differences in their spatial distributions, spectral features, and connectivity. The second stage assesses the cross-cohort consistency of these findings in a smaller, independently collected MDD cohort analyzed under the same procedure, by asking whether the effects identified in the larger cohort carry the same direction in the smaller one.

Altogether, these results further emphasize the utility of discoveries from the larger data set and the value of template-driven analyses for replicable functional network discovery from fMRI data.

### Data Sets

The primary data set we use for discovering MDD-related functional networks is the MDD-DIRECT phase II study [35]. MDD-DIRECT is a large-scale fMRI study comprised of resting-state scans from 3505 East-Asian participants across 23 data-collection sites. A subset of 1660 participants were diagnosed with Major Depressive Disorder (MDD) according to the criteria listed in the Diagnostic and Statistical Manual of Mental Disorders, 4th edition (DSM-IV). Scans from 1341 controls were collected across sites to match the age and sex ratio of MDD participants. Written informed consent was obtained from each participant, and local IRB approval was obtained at each site. Access to the data was provided in accordance with the MDD-DIRECT phase II data-sharing plan. For more details see [35].

The repetition time (TR) varies quite significantly within the full cohort (see Fig. 11); however, the majority of individuals were scanned with a TR of 2. Because sFNC and spectral effects will be significantly affected by the sampling rate, we constrain our primary analysis to a cohort of 2,427 individuals with at least a TR of 2 (1123 MDD, 831 Control, 473 indeterminate; 1207 Female, 734 Male, 486 unknown; Age 31.94+-13.93 years). Fig. 11 summarizes demographics according to diagnosis in the full population.

**Fig. 11.**
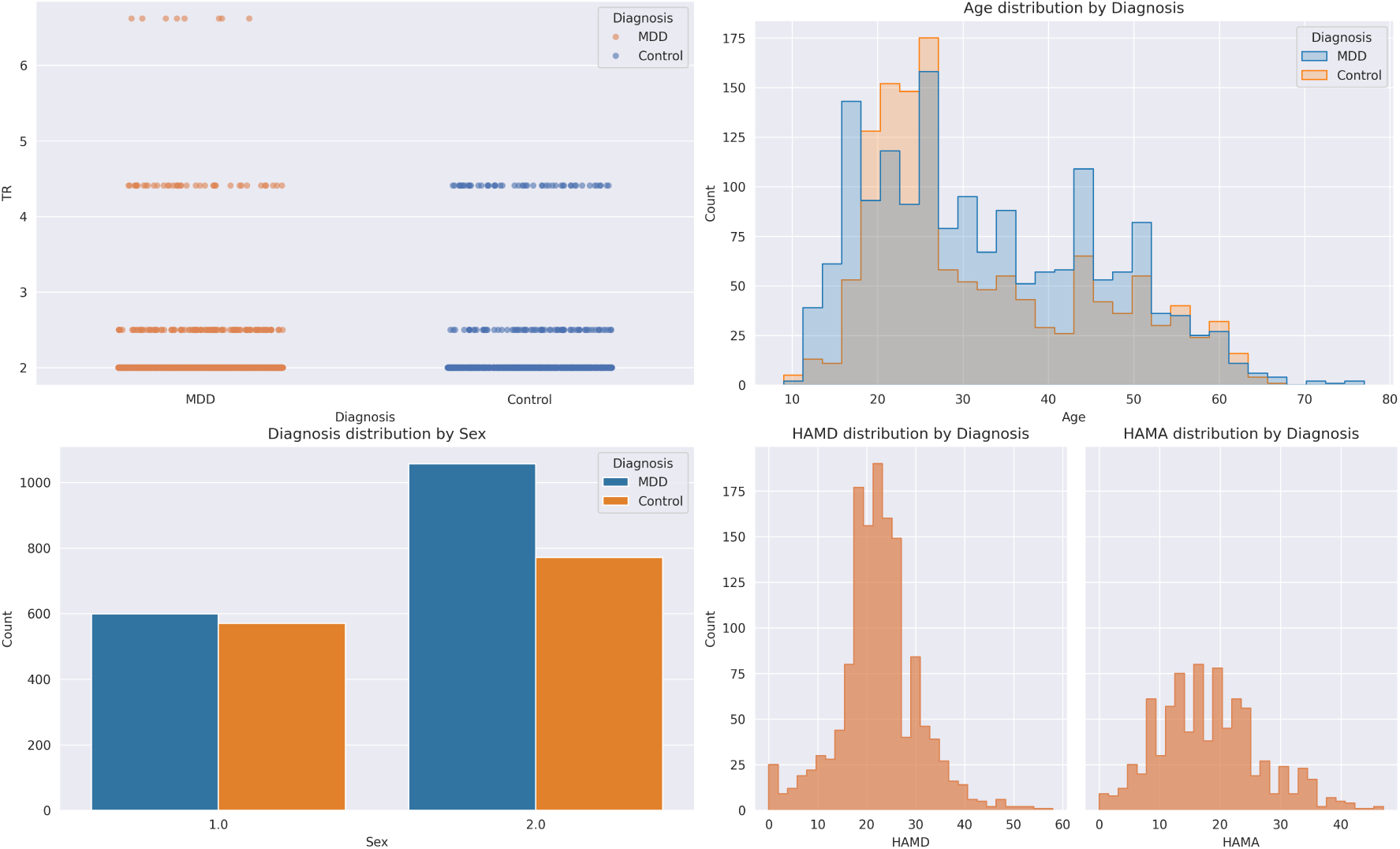
Summary of the full MDD-DIRECT cohort by diagnostic group: TR (a), age distribution (b), sex distribution (c) and symptom-severity score distribution (d). All *n* = 3,505 participants (1,660 with MDD and 1,341 controls), before restriction to the analysis cohort.

We also use a smaller, secondary data set from Zhi et al. 2018[19] to demonstrate how the networks derived from the larger MDD-DIRECT study can be applied to discovery in smaller studies. The dataset comprises fMRI scans from 519 Han Chinese participants (235 with MDD; 284 controls), recruited from four hospitals in China. All MDD diagnoses were confirmed using the DSM-IV criteria. The severity of depression symptoms was rated using the 17-item Hamilton Depression Rating Scale (HDRS). Ethical approval and informed consent were obtained, per the Institutional Review Boards at each site. For more details see [19].

### Preprocessing

Initial preprocessing was performed independently according to study protocols, with the MDD-DIRECT study using a pipeline using the Data Processing and Analysis for Brain Imaging on Surface (DPABISurf) software [38]. The functional preprocessing pipeline included reference BOLD volume generation and skull-stripping with fMRIprep[39], co-registration of the BOLD reference to the T1w image in FreeSurfer[40], head-motion parameter estimation with FSL[41], and slice-timing correction with AFNI[42]. The BOLD volumes were resampled into MNI-152 space[43, 44, 45]. Full pipeline details can be found in [35]. Additional preprocessing was performed using SPM12[46] to normalize and smooth with a Gaussian kernel at 6×6×6 FWHM, a common preprocessing step for ICA [18].

A similar preprocessing approach was adopted for the Zhi et al. 2018 data, though different software environments were used based on availability. SPM12 [46] was used in place of AFNI for slice-timing correction, for example. The same smoothing parameters were also applied to these data. As small variations in preprocessing pipelines can make knowledge transfer difficult, future studies of MDD should endeavor to use uniform preprocessing or, where possible, release raw, anonymized data to avoid this limitation.

### NeuroMark ICA

Independent Component Analysis (ICA) is a framework for solving the problem of Blind Source Separation (BSS), and has a long history of use for discovery of functional networks in both task and rest fMRI[47, 18, 48, 49, 50, 51, 52, 53]. In standard spatial ICA, the fMRI volume is modelled as composed of distinct functional networks which are independent in space, but which can covary over time. For linear ICA, the fMRI volume **X** *∈* R*^V^ ^×T^* is modelled as the linear mixture of *k* spatially independent maps **S** *∈* R*^k×V^* and their associated time-series **A** *∈* R*^k×T^* given as the product

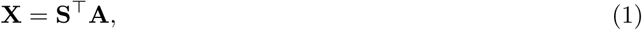

where *V* is the number of voxels, *T* is the number of timepoints, and *k* is the number of identified functional networks. These components can be obtained by optimizing a criterion for statistical independence, such as Shannon entropy in InfoMax ICA[54]. ICA can be used to discover functional networks for groups of participants by combining temporal concatenation with a post hoc back-reconstruction [18, 52].

One of the major limitations of standard “blind” ICA is that the discovery of neurobiologically-relevant functional networks requires significant manual review of estimated component maps and time-courses, especially since ICA also captures remaining noise in the data as components. Although blind ICA has been shown to reliably identify similar kinds of resting-state networks across studies [52], identifying these networks is labor-intensive and variable between studies, limiting the transfer of knowledge and ultimately impeding scientific reproducibility. NeuroMark ICA [33] provides a direct answer to these limitations of blind ICA by replacing the fully blind optimization with a spatially contained optimization, encouraging estimated components to spatially resemble to match a template of existing components. In essence, NeuroMark ICA is a bridge between data-driven and *a priori*, atlas-based functional network discovery, deriving signals directly from the data while using known signal locations to constrain the resulting networks to resemble well-studied networks such as the Default Mode, Salience, Somatomotor, and others.

Although the NeuroMark ICA framework improves ICA reliability for cross-study applications, the original version of the NeuroMark template relied on a blind ICA with a fixed model order of 100. The “best” model order for ICA of fMRI is notoriously difficult to determine, with larger model orders capturing functional networks over a broader spatial scale and smaller model orders capturing finer-grained networks. One answer to this involves performing ICA across multiple model orders to estimate functional networks of varying sizes. Recent work has involved creating a new template for NeuroMark ICA using this multi-order methodology [55], thereby expanding the available networks to include both large- and small-scale functional networks. The NeuroMark2.2 template [34] identifies 105 functional networks using this method, followed by careful manual review of derived networks in association with well-known networks identified across studies.

Using the Group ICA for fMRI Toolbox (GIFT) we apply NeuroMark ICA using the NeuroMark2.2 template to the MDD-DIRECT and Zhi et al. 2018 data sets. We perform variance normalization prior to analysis, and then perform spatially constrained ICA optimization using the default parameters in the GIFT software. Each optimization is performed independently for each scan, and the resulting spatial maps and time courses for each subject are saved prior to post-hoc analyses of spectra, spatial maps, and static functional network connectivity. Motion regression is performed with available parameters prior to the computation of sFNC and spectra.

### Static Functional Network Connectivity

Static functional network connectivity (sFNC) was computed for the resulting 105 ICA time series using NeuroMark ICA. A low-pass filter at 0.15 Hz, detrending with model order 3, and despiking were applied, all using built-in functions of the GIFT software, and according to standard methodologies for analysis of temporal features from ICA. Following postprocessing of the ICA time series, a 105 *×* 105 correlation matrix is computed such that entry in row *i*, column *j* contains the Fisher Z-scored Pearson Correlation Coefficient (PCC), providing insight into coactivation of functional networks over the full duration of the scanning period.

To study statistical differences in the sFNC, we performed cross-sectional group comparisons within each cell of the sFNC matrices from all participants with a diagnosis. We perform ANCOVA/ANOVA (and GLM regression for continuous variables) analyses at each cell, controlling for age, sex, and site as covariates and treating Diagnosis, HAMA, and HAMD scores as target variables. Local FDR correction [56] was then applied across all sFNC pairs. For the categorical diagnosis contrast, Cohen’s *d* effect sizes were computed at each FNC pair, and throughout our results we report effect sizes that are at least minor (*|d| >* 0.2). For the continuous symptom-severity scores, we report the partial correlation between score and connectivity after adjustment for covariates.

### Spectra and Frequency Analysis

**Spectra.** Time-series spectra were computed for each functional network by performing a multi-taper spectrum analysis[57] using the GIFT software. We use the GIFT defaults for this analysis, with a time-bandwidth product of 3 and *K* = 5 tapers. Within each frequency bin, we performed cross-sectional ANCOVA/ANOVA of the power in each bin for each component, again controlling for age, sex and site as covariates. Local FDR correction was applied across all frequency bins and functional networks jointly (13,545 tests).

### Spatial Maps

In a given ICA spatial map for a particular functional network, the value at a particular voxel can be understood as a continuous measure of that voxel’s contribution to the associated time series within the same component. Thus, the estimated spatial maps for each participant were also analyzed for statistical differences in the spatial distribution of each functional network. Cross-sectional statistical comparisons of the intensity values at each voxel were computed, controlling for age, sex and site as covariates. Local FDR correction was then applied across voxels and components. For the figures and voxel counts reported below, each network was restricted to its spatial extent in the NeuroMark 2.2 template, defined as the voxels where the corresponding template component exceeds *|Z| >* 2.

### Cross-Cohort Consistency

The Zhi et al. cohort was analyzed independently and identically, with local FDR applied across its own full family of tests. With 235 patients and 284 controls, a two-sided test at *α* = 0.05 has 62% power to detect an effect of *|d|* = 0.2 and 20% power for *|d|* = 0.1 even before correction for multiple comparisons, against 99% and 59% in the MDD-DIRECT analysis cohort. Since most effects in MDD-DIRECT fall in this range, the overlap of significant findings between the two cohorts is bounded largely by the power of the smaller cohort and is a poor measure of consistency. We therefore treat the smaller cohort as an assessment of cross-cohort consistency rather than as an independent replication, and use the direction of effect as the primary criterion: for each feature significant in MDD-DIRECT, we ask whether the effect in Zhi et al. has the same sign, irrespective of its significance. Agreement is compared with the 50% expected if the two cohorts were unrelated using a one-sided binomial test; because neighbouring connections, frequency bins and voxels are not independent, these *p* values are optimistic and are reported only as a guide. We also report agreement across all tests and the correlation between the unthresholded effect maps. The overlap of significant findings between cohorts is reported in Supplementary Note 2.

## Data availability

MDD-DIRECT phase II data are available to qualified researchers under the consortium’s data-sharing plan [35]. Data from Zhi et al. [19] are available from the original investigators on reasonable request.

## Code availability

NeuroMark ICA, sFNC and spectral analyses were performed with the Group ICA of fMRI Toolbox (GIFT; https://trendscenter.org/software/gift/).

## Acknowledgements

We thank the MDD-DIRECT consortium and the investigators of Zhi et al. for data access.

## Author contributions

B.B. designed the study, performed the analyses and wrote the manuscript. M.S. contributed to interpretation of results and to writing the manuscript. V.C. conceived and supervised the study and revised the manuscript.

