## Supplemental Information for "Replicable Connectivity and Spectral Alterations of Functional Networks in Major Depression"

### Supplementary Note 1: sFNC Comparisons

Supplementary Fig. 1 shows the mean sFNC matrices of the MDD and control groups in MDD-DIRECT and the difference between them.

### Supplementary Note 2: Significance Overlap Between Cohorts

This note reports the analyses of the Zhi et al. cohort on its own terms, and the overlap of significant findings between the two cohorts. Because of the limited power of the smaller cohort (Methods in the main text), these counts are a conservative and power-limited measure of consistency; the main text uses the direction of effect instead.

#### Static FNC

We next investigated whether these connectivity differences are recovered in the smaller, independently collected cohort of Zhi et al. [1]. Analyzed independently and identically, with local FDR applied across its own full set of 5460 tests, that cohort yields 307 significant connections (5.6%), 161 increased and 146 decreased in MDD. **125 connections are identified independently in both studies.** If the two sets were unrelated, an overlap of approximately 37 connections would be expected by chance ($651 \times 307/5460$), so the observed overlap is more than three times the chance rate. All 125 agree in the direction of effect. The connectivity differences in this cohort are shown in Supplementary Fig. 2.

The connections that replicate are concentrated in the visual-to-temporal and default-mode blocks. The largest replicated groups are VI-OC–HC-TP (18 connections, all reduced; mean $d = -0.226$ in MDD-DIRECT and $-0.266$ in Zhi et al.), CB–SM (7, 6 increased), VI-OT–HC-TP (7, all reduced), PL–TN-DM (7, all increased), HC-FR–TN-DM (7, all increased) and HC-FR–SM (5, all reduced). The most frequently replicated individual networks are posterior superior temporal gyrus (11 connections, 10 reduced), superior sensorimotor cortex (10, 8 increased), dorsal precuneus (8) and bilateral temporoparietal junction (7, all reduced). The single best-replicated connection is also the largest effect in the primary analysis: occipital pole to dorsal precuneus, $d = -0.403$ in MDD-DIRECT and $d = -0.374$ in Zhi et al. ($p = 4.3 \times 10^{-11}$ and $5.4 \times 10^{-5}$). By contrast the cerebello-striatal and cerebello-thalamic blocks, the densest findings in MDD-DIRECT, contribute no co-identified connections. Co-identified effects were of comparable or slightly larger magnitude in the smaller cohort (mean $|d| \approx 0.19$ versus 0.24); because these connections were selected in part on their significance in the smaller cohort, this difference is expected on statistical grounds and should not be read as a stronger effect.

#### Spectra

Analyzed independently under local FDR across its own full family of 13,545 tests, the Zhi et al. cohort yields 625 significant bins (4.6%). **59 bins are identified independently in both studies**, against approximately 20 expected by chance ($432 \times 625/13{,}545$), and all 59 agree in direction. The frequency dependence itself is reproduced, with a correlation of 0.41 between frequency and effect direction among the

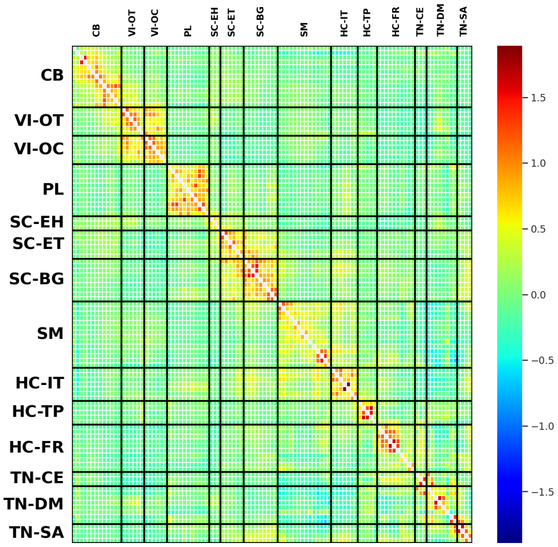


**a** Mean sFNC matrix, MDD group.

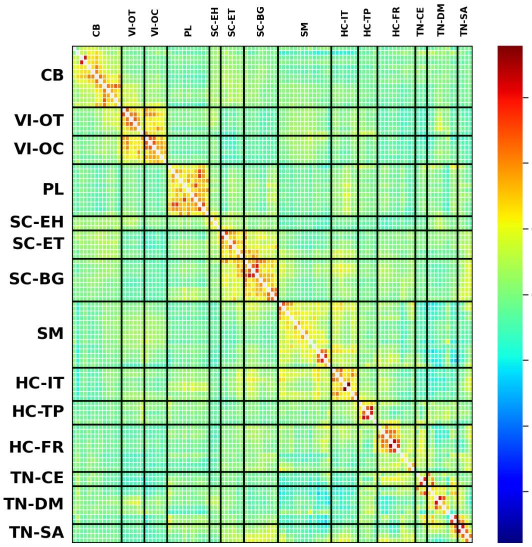


**b** Mean sFNC matrix, control group.

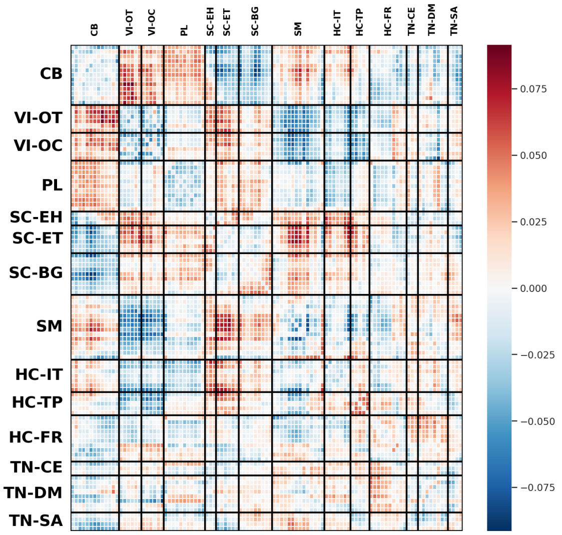


**c** Difference between the group mean sFNC matrices (MDD minus control).

**Supplementary Fig. 1 | Group mean sFNC matrices and their difference.** Values are the mean Fisher $z$-transformed correlation across participants in MDD-DIRECT ($n$ = 1,123 patients with MDD and 831 controls).

59 co-significant bins. The largest single replicated group is in posterior cingulate cortex (11 bins, all showing increased power in MDD, $d = +0.189$ rising to $+0.223$), a network that contributes comparatively little to the primary analysis; the most precisely reproduced effect is the right thalamus reduction at 0.008–0.016 Hz, which appears in both cohorts in the same narrow band, in the same direction and at comparable magnitude ($d = -0.186$ and $-0.222$). All 59 replicated bins lie at or below 0.162 Hz. The spectral differences in this cohort are shown in Supplementary Fig. 3.

### Spatial Maps

Analyzed independently under the same criteria, the Zhi et al. cohort yields a far larger number of surviving voxels than MDD-DIRECT (194,075 at $|d| \geq 0.2$), but this reflects the behaviour of standardized effect sizes

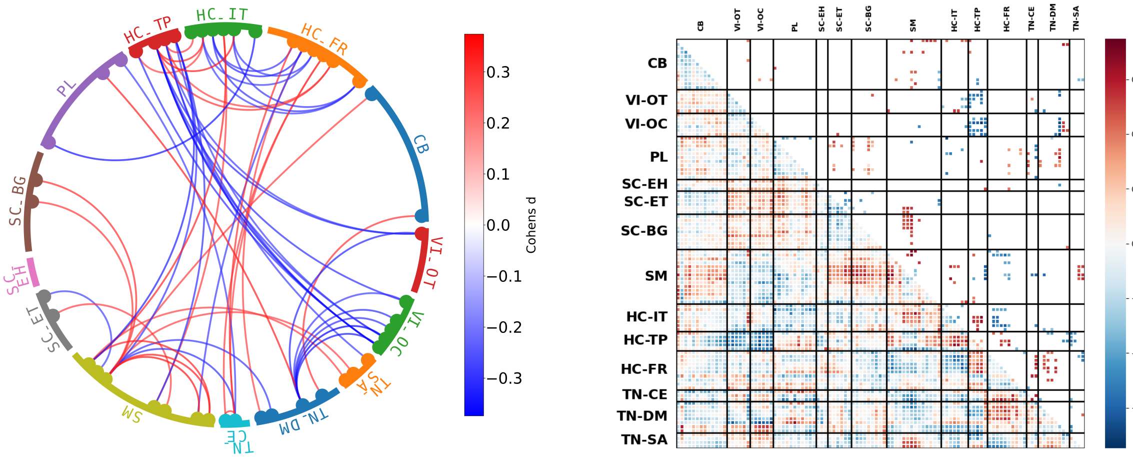


**a** Chord diagram of the top 55 connections ( 1% of 5460) in the Zhi et al. cohort, ranked by Cohen's $d$, among those significant in that cohort's own independent local-FDR analysis. Colour indicates Cohen's $d$; warm colours denote higher connectivity in MDD and cool colours lower connectivity in MDD.

**b** Full sFNC effect-size matrix for the Zhi et al. diagnostic contrast, analyzed independently. Lower triangle: all connections. Upper triangle: connections significant after local FDR within this cohort.

**Supplementary Fig. 2 | Static functional network connectivity differences between MDD patients and controls in the Zhi et al. cohort, analyzed independently.** $n = 235$ patients with MDD and 284 controls. Each connection was compared between groups by ANCOVA adjusting for age, sex and site, with local FDR correction ($q \leq 0.05$) across all 5,460 connections in this cohort; effect size is Cohen's $d$.

in smaller samples rather than a stronger effect. The distribution of $|d|$ in that cohort is correspondingly wider (median 0.060 against 0.037; maximum 0.502 against 0.319), as expected at $N = 519$. Unlike MDD-DIRECT, however, the surviving voxels are directionally balanced within networks (mean within-network sign consistency 0.535 against 0.714 in MDD-DIRECT), so that the coherent gains and losses of voxel loading that characterize the primary analysis are not recovered. On the same independent-analysis criterion applied to connectivity and spectra, the spatial-map differences therefore do not reproduce. The spatial-map differences in this cohort are shown in Supplementary Fig. 4.

### Symptom Severity

In the Zhi et al. cohort, 179 connections were associated with HDRS total score, 80 positively and 99 negatively. Only three connections were co-identified across studies. Associations between spatial-map intensity and HDRS total score in this cohort are shown in Supplementary Fig. 5.

### Discussion of Significance Overlap

We have deliberately assessed replication in a conservative way. The Zhi et al. cohort was analyzed independently and identically, with local FDR applied across its own full family of tests, and we report the intersection of the two resulting sets of findings. We did not restrict testing in the replication cohort to the features identified in MDD-DIRECT. That alternative is common in the literature and would have produced considerably more favourable numbers, but the concordance it yields is partly a property of the selection rather than of the replication data, and it cannot distinguish a genuinely reproduced effect from one that merely fails to be contradicted in a smaller sample.

On this stricter criterion, connectivity and spectra both reproduce. 125 connections and 59 spectral bins are identified independently in both cohorts, against chance expectations of approximately 37 and

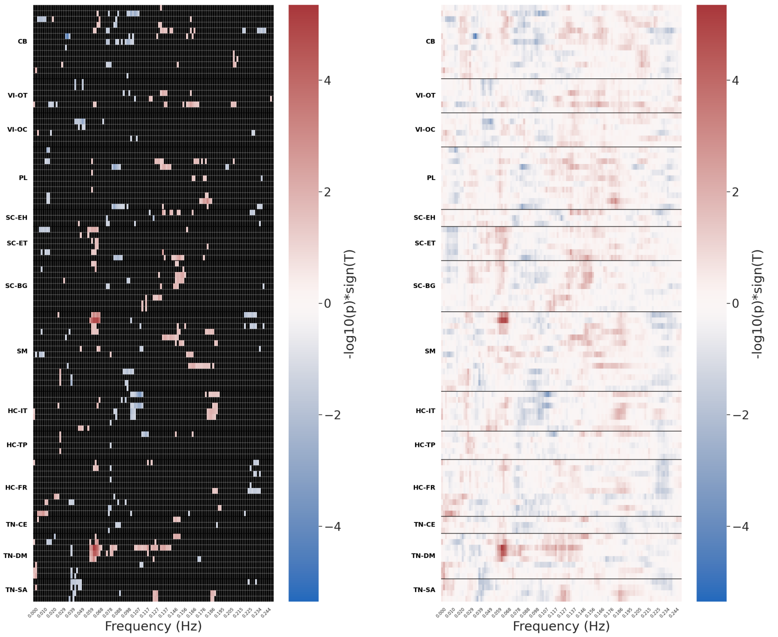


**a** Spectral power differences in the Zhi et al. cohort, restricted to bins significant in that cohort's own independent local-FDR analysis.

**b** The same comparison without thresholding.

**Supplementary Fig. 3 | Spectral power differences between MDD patients and controls in the Zhi et al. cohort, analyzed independently.** $n = 235$ patients with MDD and 284 controls. Power in each of 129 frequency bins of each of 105 networks was compared between groups by ANCOVA adjusting for age, sex and site, with local FDR correction ($q \leq 0.05$) across all 13,545 network–frequency tests in this cohort; effect size is Cohen's $d$.

20 respectively, and in both cases every co-identified effect agrees in direction **across all connections**. Directional agreement at that rate is not guaranteed by the overlap itself, since two independently significant sets could in principle overlap while disagreeing in sign, and it is the aspect of the result we find most persuasive. The spatial maps, analyzed the same way, do not reproduce.

One asymmetry in the replication effort deserves further comment. The cerebello-subcortical block is the densest finding in MDD-DIRECT but contributes no replicated connections, while the visual-to-temporal and cerebello-somatomotor blocks replicate well. Differences in cerebellar coverage between the two acquisitions are a plausible contributor, as are susceptibility and normalization differences affecting the caudate and striatal components most strongly. Their direction of effect, however, agrees in 75 of 81 connections (Discussion in the main text), indicating that the absence of overlap reflects the power of the smaller cohort rather than inconsistency between cohorts. The spectral results show a comparable pattern in reverse: posterior cingulate cortex contributes comparatively little to the primary analysis but the largest single group of reproduced bins, indicating that prominence within one cohort is a poor guide to reproducibility across cohorts.

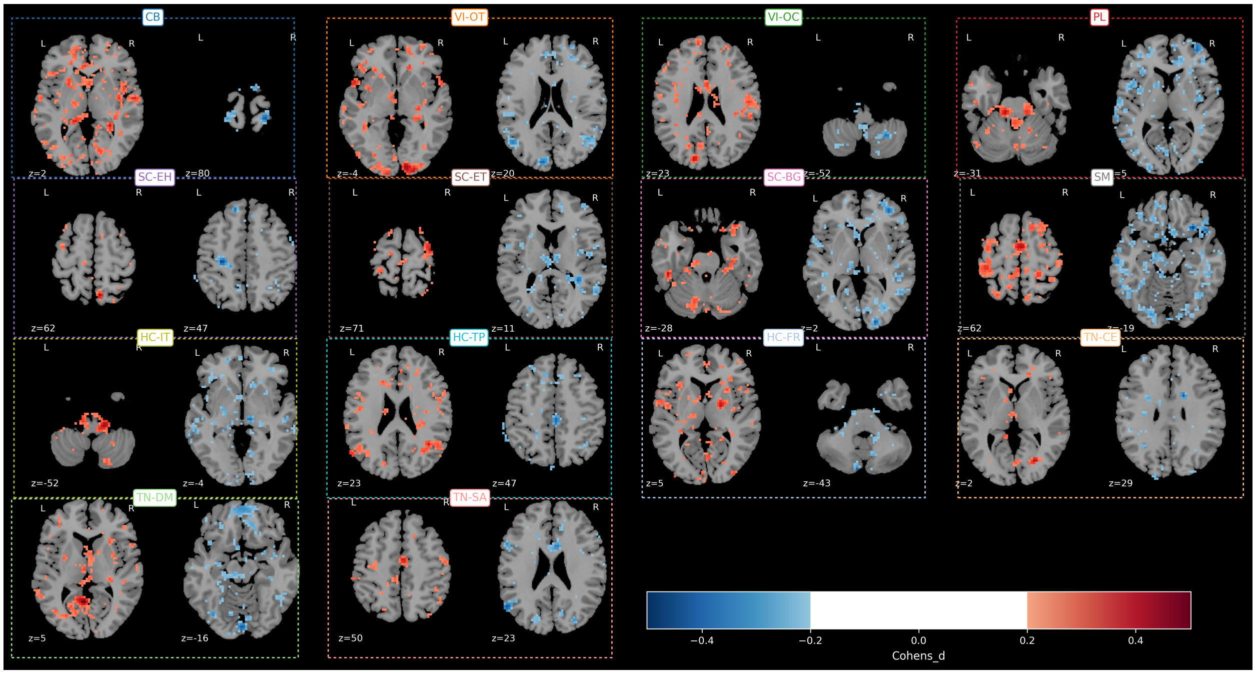


**a** Networks showing the largest positive and negative differences in voxel loading between MDD patients and controls, by domain.

**Supplementary Fig. 4 | Spatial map differences between MDD patients and controls in the Zhi et al. cohort.** Positive values denote higher voxel loading on that network in MDD. $n = 235$ patients with MDD and 284 controls. Voxel loadings were compared between groups by ANCOVA adjusting for age, sex and site. Voxels are shown where significant after local FDR ($q \leq 0.05$) with $|d| \geq 0.2$; colour shows Cohen's $d$. Unlike Fig. 5, voxels are not restricted to the template extents.

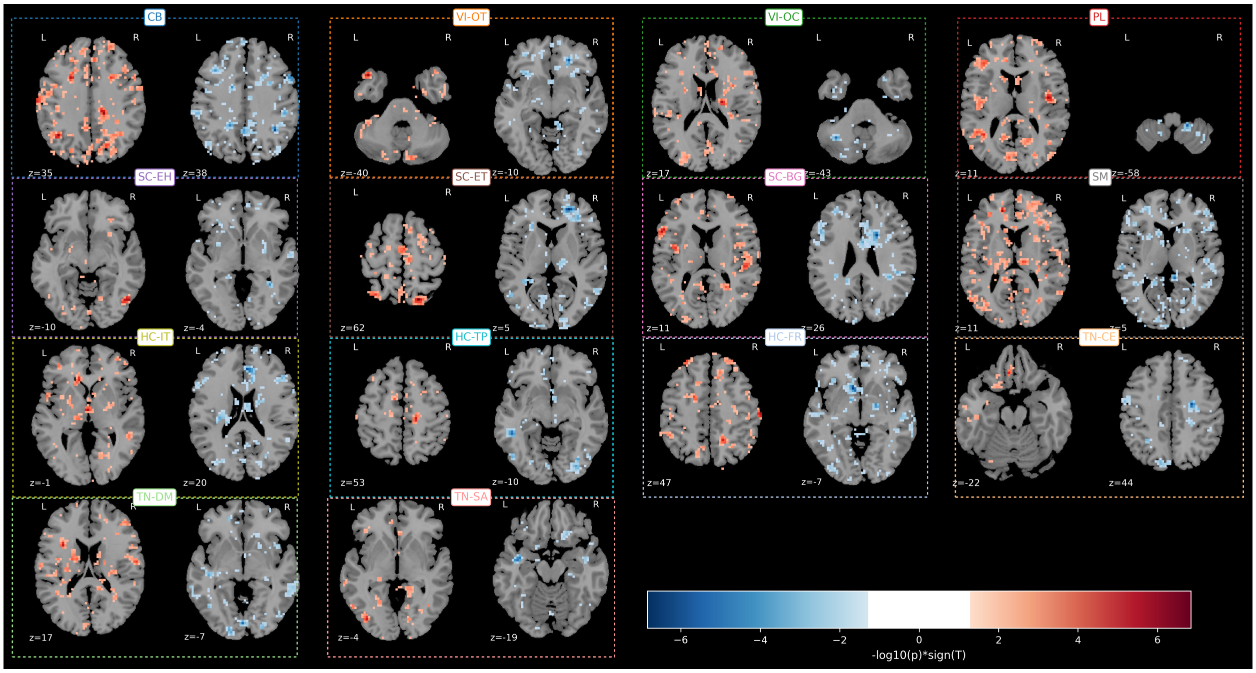


**a** Networks showing the largest positive and negative associations between voxel loading and HDRS total score, by domain.

**Supplementary Fig. 5 | Associations between spatial map intensity and HDRS score in Zhi et al.** Voxel loadings were regressed on HDRS total score, adjusting for age, sex and site.
